# Increased capacity to maintain glucose homeostasis in a transgenic mouse expressing human but not mouse growth hormone with developing high fat diet-related insulin resistance, steatosis and adipose dysfunction

**DOI:** 10.1101/2024.02.21.581281

**Authors:** Yan Jin, Jessica S Jarmasz, Shakila Sultana, Luis Cordero-Monroy, Carla G Taylor, Peter Zahradka, Elissavet Kardami, Peter A Cattini

## Abstract

**Objective:** Differences in primate and non-primate growth hormone (GH) genes can affect their regulation and function. This includes the ability of human (h) but not mouse (m) GH to bind the prolactin (PRL) receptor (PRLR) as well as the GH receptor (GHR). Potential differential effects were assessed in male hGH- or mGH-expressing mice and fed high fat diet (HFD) *versus* regular chow diet (RCD). Pancreas and epididymal white adipose tissue (eWAT) gene expression and/or related function were targeted as the pancreas responds to both PRLR and GHR signaling and catabolic effects like lipolytic activity are more directly attributable to GH and GHR signaling.

**Design:** A transgenic CD-1 mouse expressing hGH but not mGH under hypothalamic control was generated to compare with wild type CD-1 mice and size and bone density determined. Glucose clearance, islet area, insulin and insulin-like growth factor (IGF) -2 gene expression were assessed as well as serum glucose and insulin levels in mice fed a HFD *versus* RCD for 8 and 24 weeks. Adiposity, liver and serum triglycerides as well as eWAT cell area, cytokine (leptin and adiponectin) and senescence-related marker (p21^CIP1^ and p16^INK4a^) RNA levels were also assessed.

**Results:** Male hGH-expressing transgenic CD-1[ΔmGH.hGH] mice have significantly lower liver IGF-1 RNA levels and are smaller based on length and weight than wild type CD-1[mGH] mice. They also have ∼1.5-fold higher total body fat and serum triglyceride levels. However, CD-1[ΔmGH.hGH] and CD-1[mGH] mice grow at the same rate with similar cortical and trabecular bone densities. Unlike CD-1[mGH] mice, there was no significant delay in glucose clearance in CD-1[ΔmGH.hGH] mice after 8 weeks on a HFD *versus* RCD; while basal (RCD) serum insulin levels were similar, fasting glucose levels were lower and pancreas IGF-2 RNA levels were increased in CD-1[ΔmGH.hGH] mice. However, both CD-1[ΔmGH.hGH] and CD-1[mGH] showed evidence of increased insulin resistance after 24 weeks on HFD, including delayed glucose clearance in spite of increased pancreatic islet area and insulin production as well as signs of liver steatosis and increased hepatic triglyceride levels. These increases correlated with elevated PRLR but not GHR RNA levels. Assessment of eWAT revealed >2-fold larger adipocytes in CD-1[ΔmGH.hGH] compared to CD-1 [mGH] mice fed RCD at both 12 and 28 weeks. This was associated with an ∼2.6-fold increase in leptin RNA levels at 12 weeks and ∼58% lower adiponectin RNA levels at 28 weeks. A >2-fold increase in p21^CIP1^ transcript levels was also detected in eWAT from both CD-1[ΔmGH.hGH] and CD-1 [mGH] mice fed RCD with age (28 *versus* 12 weeks) but were unaffected by diet. However, a >2-fold increase in p16^INK4a^ RNA levels was observed after 24 weeks on HFD.

**Conclusions:** While limited to observations in the male, transgenic CD-1[ΔmGH.hGH] mice exhibit signs of GH insufficiency and eWAT adipocyte dysfunction. These mice also show an initial resistance to the negative effects of HFD on glucose clearance when compared to CD-1[mGH] mice, which is potentially related to a differential effect of hGH *versus* mGH on pancreas development and/or function.

## INTRODUCTION

Growth hormone (GH) is produced and secreted specifically by the somatotrophs of the anterior pituitary gland [1, 2]. Pituitary GH is generally regarded as a requirement for regulating somatic growth in vertebrates, including rodents and primates [3, 4], but like other structurally related cytokines in humans, plays an important role controlling health and disease processes [5]. GH can exert its effects directly through the GH receptor (GHR) or indirectly through the stimulation of insulin-like growth factor-1 (IGF-1) [6, 7]. GH reaches its highest circulatory levels in puberty, where it contributes to energy homeostasis and somatogenic growth [8]. Unlike the anabolic effects of GH on many tissues, the catabolic lipolytic effect on adipose tissue is mediated largely by GH and its receptor, and not the actions of IGF-1 or insulin [7, 9]. In this context, adipose tissue represents a potentially useful target when assessing GH activities including possible differential effects of GHs and, specifically, of primate and non-primate origin, since human (h) GH and rodent GH genes differ significantly in structure and thus have the capacity for different regulation, receptor binding, activities and functions [10, 11]. In the pancreas, insulin-producing β-cells represent another useful target to assess GH-related activity. Rodent and other non-primate GHs are unable to bind the prolactin (PRL) receptor (PRLR), while primate GHs, including hGH, can associate with non-primate GH receptor (GHR) and PRLR with high affinity [10]. Thus, hGH, unlike mouse (m) GH possesses lactogenic as well as somatotropic activity, which may have implications in terms of the pancreas, since GHR and to a greater extent PRLR activation have been linked to effects on pancreatic β-cell mass [12–16]. Furthermore, IGF-2 levels in the pancreas are more strongly linked to PRLR signaling (via PRL and placental lactogen) *versus* the GHR [17, 18], and in the adult pancreas IGF-2 may act in an autocrine manner to control β-cell mass and/or help maintain glucose-stimulated insulin secretion in aging, pregnancy, metabolic stress and with induction of insulin resistance [19, 20].

Multiple adipose tissue depots exist and these have been described broadly as either subcutaneous (located beneath the skin) or intra-abdominal based on location [7]. While subcutaneous white adipose tissue (WAT) has not been associated with disease risk, visceral WAT that is distributed around organs is often associated with higher risk for type 2 *diabetes mellitus*, hypertension and all-cause mortality [7, 21]. Although there are species-specific differences in depot distribution [7], the perigonadal (epididymal or paraovarian) depot is most commonly studied as WAT in mice due to its accessibility and mass [7], and thus epididymal WAT (eWAT) was also targeted for study here. WAT is more abundant than brown adipose tissue and white adipocytes can make up to 90% of WAT volume, although intermediary adipocytes may be present [7]. WAT has endocrine activity and secretes multiple adipokines including leptin and adiponectin [7]. Leptin is a satiety hormone while adiponectin is an anti-inflammatory and insulin-sensitizing hormone [7]. Whether in patients or mice, a decrease in GH signaling is associated with an increase in both hormone levels [9], and modified leptin and adiponectin gene expression is an indicator of adipose tissue dysfunction. More specifically, leptin RNA levels correlate with adipocyte volume in each fat depot in mice and, thus, are expected to increase with obesity [22, 23], while adiponectin transcript levels are reduced in obesity and in insulin resistant states in mice and humans [24, 25].

Here, we attempted to gain an insight into possible differential effects of hGH and mGH in an *in vivo* context in post-pubertal male mice, including chronic (8 and 24 weeks) high fat diet (HFD) on the pancreas and eWAT. A transgenic mouse expressing hGH, but not mGH, in a pituitary-specific manner and under hypothalamic control was generated, and effects on adiposity, steatosis and glucose clearance were assessed. In addition, RNA levels for pancreatic MAF bZIP transcription factor A (MAFA), insulin (Ins1 and Ins2), IGF-2, GHR and PRLR as well as white adipose tissue (WAT) leptin, adiponectin peroxisome proliferator-activated receptor γ (PPARγ; adipogenesis and lipid storage-related) [9, 26], and senescence markers p21^CIP1^ and p16^INK4a^, were assessed in mice fed a high fat diet (HFD) compared to a regular chow diet (RCD) for up to 24 weeks. Previously, transgenic CD-1[mGH.hGH] mice with a single copy of the hGH gene locus in addition to the endogenous mGH gene were described [11, 27]. The transgene includes the hGH locus control region required for pituitary-specific expression *in vivo* and, as a result, these mice have been used to study the response of the hGH gene to hormone treatments [28, 29], diet [30], light/day cycle [31, 32] and sleep deprivation [33] in a physiological context. Here, we have removed the endogenous mGH gene from CD-1[mGH.hGH] mice by crossing it with the Ghtm1(KOMP)Vlcg mouse (UCDAVIS KOMP repository; also referred to as C57BL/6[Gh^tm1Vlcg^] or GH^−/−^ mouse [34]) to generate CD-1[ΔmGH.hGH] mice, which allow the effects of hGH, without mGH, to be assessed *in vivo*. CD-1[ΔmGH.hGH] mice were then maintained on a CD-1 mouse genetic background for ten generations before initial characterization of mouse size, hGH levels, response to a GH releasing peptide (GHRP-2) and bone density in mice were evaluated. Subsequently, the effect of chronic (8 and/or 24 weeks) HFD on glucose clearance, pancreatic insulin, IGF-2, GHR and PRLR gene expression, serum triglycerides and liver steatosis were examined, as well as eWAT adipocyte area and cytokine (leptin and adiponectin) and senescence-related marker (p21^CIP1^ and p16^INK4a^) transcript levels.

While limited to a study of male mice, our observations provide evidence of GH insufficiency and eWAT adipocyte dysfunction in CD-1[ΔmGH.hGH] compared to wild type CD-1[mGH] mice. However, while both hGH- and mGH-expressing mice showed signs of developing insulin resistance on 8 weeks HFD in terms of liver discoloration and triglyceride content, glucose clearance was impaired in wild type CD-1[mGH] but not transgenic CD-1[ΔmGH.hGH] mice, which express higher relative levels of IGF-2. This differential glucose clearance-related response was no longer seen after a more prolonged time (24 weeks) on HFD, when a greater increase in insulin resistance in both hGH and mGH-producing mouse types was suggested by effects on pancreatic β-cells and insulin gene expression. These observations are discussed in relation to GH levels and the ability of hGH to potentially activate both somatogenic and lactogenic pathways.

## MATERIALS AND METHODS

### Mice, diet and breeding

All procedures involving animals and their tissues conform to the Guide for the Care and Use of Laboratory Animals published by the Canadian Council on Animal Care (https://ccac.ca/Documents/Standards/Guidelines/Experimental_Animals_Vol1.pdf) and were done with approval of the Animal Protocol Management and Review Committee at the University of Manitoba. To generate CD-1[ΔmGH.hGH] mice, the CD-1[mGH.hGH] mouse (also referred to as F-74, 171 hGH/CS or hGH/PL) that contain a single copy of the hGH gene and locus control region [27] were crossed with the C57BL/6[Gh^tm1Vlcg^] mouse, which is also known as the GH^−/−^ mouse [34]. The mGH gene coding region on chromosome 11 has been deleted in this mouse and replaced with the β-galactosidase gene (*LacZ*) as described elsewhere (http://velocigene.com/komp/detail/10368) [34]. A program of 10 generations of breeding was followed to clear the C57BL/6 mouse genetic background as well as generate a homozygous line. A total of 1,381 pups were derived from 210 pregnancies, suggesting an average litter size of 6-7 pups, which contrasts with ∼12 reported for wild type CD-1[mGH] mice [35]. All CD-1 mouse types were housed in an environmentally controlled room maintained on a 12-hour light/dark cycle (6 a.m./6 p.m.). Access to food in the form of palatable pellets and tap water was *ad libitum*.

For studies involving a challenge with high fat diet, male mice were fed a regular chow diet (RCD; fat 14 kcal%, carbohydrate 60 kcal% and protein 25 kcal%; <u>Pro-lab, RMH3000 5P00</u>) but at postnatal week 4, mice were randomly assigned to a high-fat diet (HFD; fat 60 kcal%, carbohydrate 20 kcal% and protein 20 kcal%; <u>Research Diets, D12492</u>) or maintained on the RCD for 8 weeks or 24 weeks. Therefore, all RCD fed mice were on the diet for 12 and 28 weeks while HFD fed mice were on RCD for the first 4 weeks and then switched to HFD for 8 and 24 weeks. Fat content in both diets is lard and soybean oil.

To stimulate and measure GH secretion, 4 week-old CD-1[ΔmGH.hGH] mice fed RCD, were switched to either HFD or maintained on RCD for 4 weeks (4 mice per diet group). Mice were given a single 330 μg/kg dose of a synthetic growth hormone releasing peptide-2 (GHRP-2; #OPPA01036, Aviva Systems Biology) by intra-peritoneal (i.p.) injection, which acts like GH releasing hormone [36]. Blood was taken from the saphenous vein, before and 10 minutes post-injection for assessment by enzyme-linked immunosorbent assay (ELISA) of serum separated by centrifugation (9,300 g for 10 minutes) at 4 °C.

### Glucose tolerance test (GTT)

A GTT was performed on adult mice at 12 and 28 weeks essentially as described previously [37]. Briefly, all mice were weighed, fasted for approximately 16 hours and then a GTT was performed using 2 g/kg of intraperitoneal glucose (Sigma, G7258) dissolved in double-distilled water. Blood was collected from a tail vein and glucose levels (mM) were assessed prior to injection (0 minutes), and then 15, 30, 60, 90 and 120 minutes after the glucose injection using a glucose meter (OneTouch Ultra2 Glucose Monitoring System, Johnson & Johnson, Lifescan, Inc) [38].

### Bone density and whole-body fat scans

Computed tomography (CT) scans were conducted on a Skyscan 1776 micro-CT scanner (Bruker, Kontich, Belgium) in the Central Animal Core Imaging and Transgenic Facilities at the University of Manitoba (https://umanitoba.ca/health-sciences/research/central-animal-core-imaging-and-transgenic-facilities). For assessment of bone density and whole body fat, 4 week-old CD-1[ΔmGH.hGH] and CD-1[mGH] mice fed RCD, were switched to either HFD or maintained on RCD for 24 weeks (8 mice per diet group). These mice were anesthetized with 4% isoflurane in oxygen and maintained with 2-2.5% isoflurane in oxygen before bone density and body fat scanning. CT scans of the femur were acquired with 9 μm spatial resolution, a 0.5 mm Al filter, source voltage and current of 50 kV and 500 μA, respectively, and 1,175 seconds acquisition time. Whole-body fat scans were acquired with a spatial resolution of 35 μm, a 1 mm Al filter, source voltage and current of 65 kV and 385 μA, respectively, and 80 second acquisition time.

Shadow projections were reconstructed into 3-dimensional CT images using the Nrecon software package (Bruker, Kontich, Belgium). Bone scans were reconstructed using a ring artefact correction factor of 4, a beam hardening factor of 30% and dynamic range of 0-0.1. Whole-body fat scans were reconstructed using a ring artefact correction factor of 4, a beam hardening factor of 30% and variable dynamic range. Bone density and whole body fat percentage were calculated using micro-CT software (CTAn, Bruker, Kontich, Belgium).

Cortical bone density was measured from 100 slices (900 μm) and 250 slices (2,250 μm) from the growth plate. Trabecular bone density was calculated from a 200 slice (1,800 μm) region starting 50 slices (450 μm) from the growth plate. Results for bone density are given in grams hydroxyapatite/cm^3^ (g/cm^3^). Whole-body fat values are calculated as volume of fat/body volume x100.

### Serum biochemistry and hepatic triglycerides

At termination of the study, serum was prepared from non-fasted blood and analyzed for triglycerides, LDL-cholesterol, HDL-cholesterol and total cholesterol on an Autoanalyzer c111 (Roche Diagnostics). Liver weight was also measured and liver homogenates were assayed for triglycerides (LabAssay™ Triglyceride colorimetric assay; FUJIFILM Wako Pure Chemical Corporation).

### Estimation of liver discoloration (fatty liver/steatosis)

Liver discoloration was used to estimate steatosis in 12 and 28 week-old mice after 8 and 24 weeks, respectively, on HFD. Briefly, liver images were captured digitally and then assessed in a blinded-study using a numerical six point (0-5) scale reflecting increasing level of liver discoloration due to fat content. The scores from each mouse and diet group were averaged and a score of 1 or less represents a relatively healthy liver and a score of greater than 3 reflects a relatively unhealthy liver (evidence of steatosis).

### Detection of GH and insulin by ELISA

Blood was taken from the saphenous vein (∼0.15 ml) and serum separated by centrifugation (9300 g for 10 minutes) at 4 °C. Serum samples were stored at −80 °C until assessed by ELISA according to manufacturer’s instructions (Ultrasensitive Human Growth Hormone ELISA, 22-HGHHUU-E01, ALPCO; mGH, EZRMGH-45K and Rat/Mouse Insulin ELISA, EZRMI-13K, Millipore).

### Real-time reverse transcription polymerase chain reaction (PCR)

Tissues (anterior pituitary, liver, pancreas and epididymal WAT (eWAT)) were collected and flash-frozen after euthanasia by cervical dislocation before isolation of total RNA. Pituitary RNA was extracted using the RNeasy Plus Mini Kit (Qiagen) with Qiagen QIA Shredder™ 250. Liver, pancreas and eWAT RNA was extracted using the RNeasy Plus Universal Mini Kit (Qiagen). RNA (1 μg) was reverse transcribed using the QuantiTect Reverse Transcription Kit (Qiagen) according to the manufacturer’s instructions. Quantitative PCR (qPCR) and/or PCR were done using specific primers for pituitary hGH, mGH, liver IGF-1, pancreas GHR, IGF-2, Ins1, Ins2, MAFA and PRLR as well as eWAT adiponectin, leptin, PPARγ, p16^INK4a^, p21^CIP1^ and glyceraldehyde-3-phosphate dehydrogenase (GAPDH) gene transcripts, with primer sequences as described (Table 1). Gene expression level in each sample (absolute quantification) was calculated from a standard curve and normalized to mouse GAPDH expression as appropriate. Tests were run in duplicate on 4-12 independent samples. For functional hGH and mGH gene detection in mouse pituitary tissue by PCR, hGH and mGH amplicons were assessed by 1.5% agarose gel electrophoresis and ethidium bromide staining.

**Table 1:**
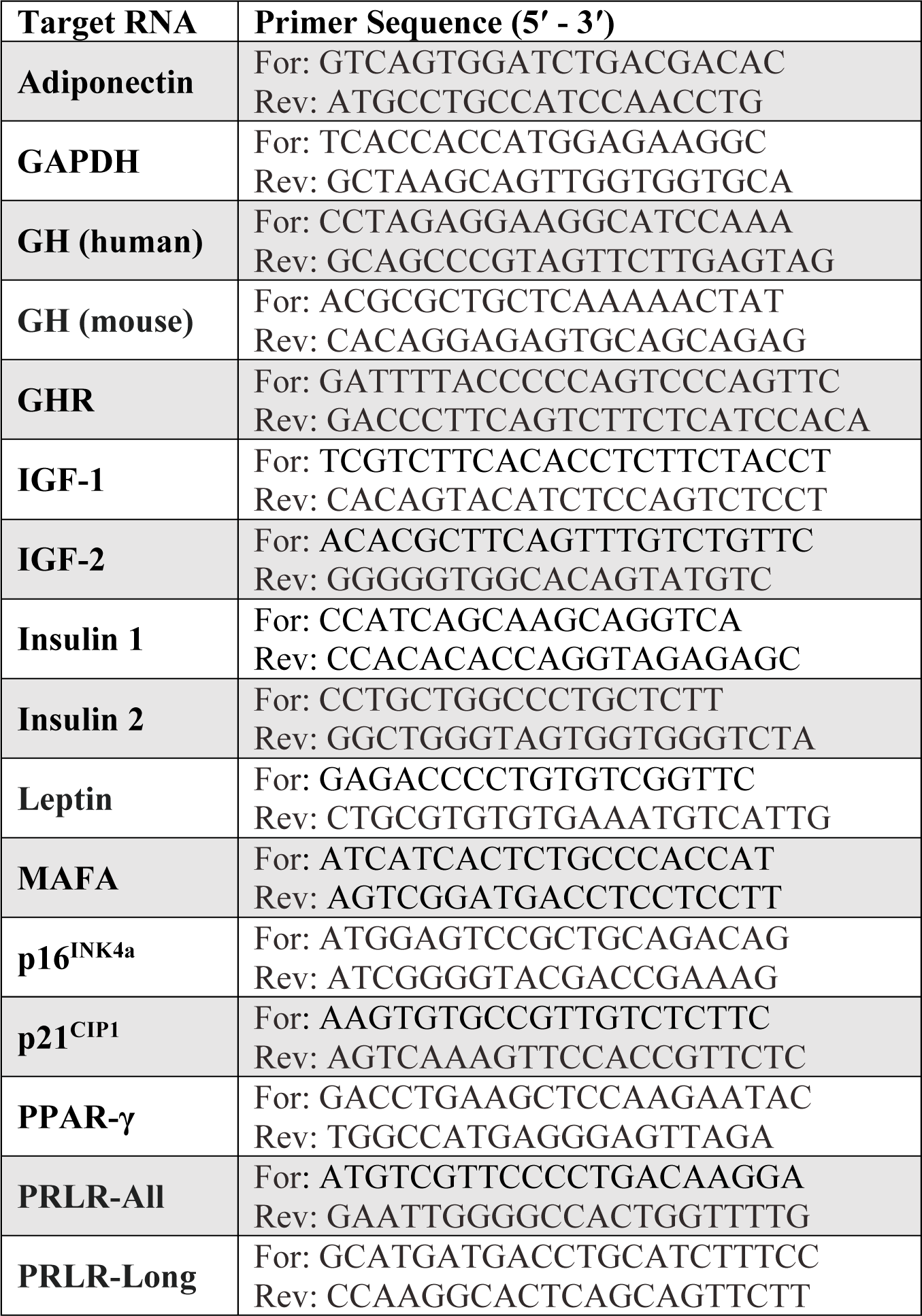
Forward (For) and reverse (Rev) primers used for polymerase chain reaction.

### Histology and determination of adipocyte and pancreatic islet areas

Epididymal WAT and pancreas were fixed in 10% formalin in phosphate-buffered saline for 24 hours, and then paraffin-embedding, sectioning (5-μm sections) and staining of nuclei and cytoplasm with hematoxylin and eosin (H&E) were done through the Histology Services, Imaging Facility, Electron Microscopy Platform at the University of Manitoba (https://umanitoba.ca/health-sciences/research/histology-services-imaging-facility-electron-microscopy-platform) while H&E staining of pancreas was completed in the laboratory. Images of slides were captured with an IX81 Inverted Fluorescence Microscope (Olympus, Centre Valley, PA, USA) using Infinity Analyze Software (Lumenera, Ottawa, ON, Canada). As a measure of size, adipocyte area and pancreatic islet area were quantified with the open-source image analysis program ImageJ 1.52 (ImageJ, US Nation Institutes of Health, Bethesda, MD, USA) [39]. A continuous block of an equal number of adipocytes from each available adipose section for a total of 40 adipocytes per animal was used to determine average cell area (μm^2^) for each treatment group [40]. Pancreatic islet area (μm^2^) was analyzed using three separate tissue sections sufficiently spaced for visualization of unique islets.

### Statistical analysis

Prism 9.5.1 software was used for statistical analysis. Three-way or two-way ANOVA with Tukey’s *post hoc* test were used for multiple comparisons as appropriate (combination of mouse type and/or diet and/or assessment time). In figures, where three-way ANOVA-related data are presented, significant differences are identified between an effects of diet within a mouse type (*), an effect of time on a single diet within a mouse type (‡) and the effect of a specific diet between mouse types (#). A value of p<0.05 was considered statistically significant and represented in figures as: */#/‡, p<0.05; **/##/‡‡, p<0.01; ***/###/‡‡‡, p<0.001; ****/####/‡‡‡‡, p<0.0001. All error bars represent the standard error of the mean (SEM).

## RESULTS

### GH expression, body weight, response to GHRP-2 injection and liver IGF-1 RNA levels

Total RNA was isolated from the pituitary gland of CD-1[mGH], CD-1[mGH.hGH] and CD-1[ΔmGH.hGH] mice at 8 weeks of age who were on a RCD, and assessed for GH using specific primers for hGH and mGH transcripts by PCR. As expected, only hGH RNA was detected in the pituitary of CD-1[ΔmGH.hGH] mice (Figure 1). Both hGH and mGH transcripts were detected in CD-1[mGH.hGH] mice and mGH RNA alone was detected in the wild type CD-1[mGH] mouse pituitary sample (Figure 1).

**Figure 1.**
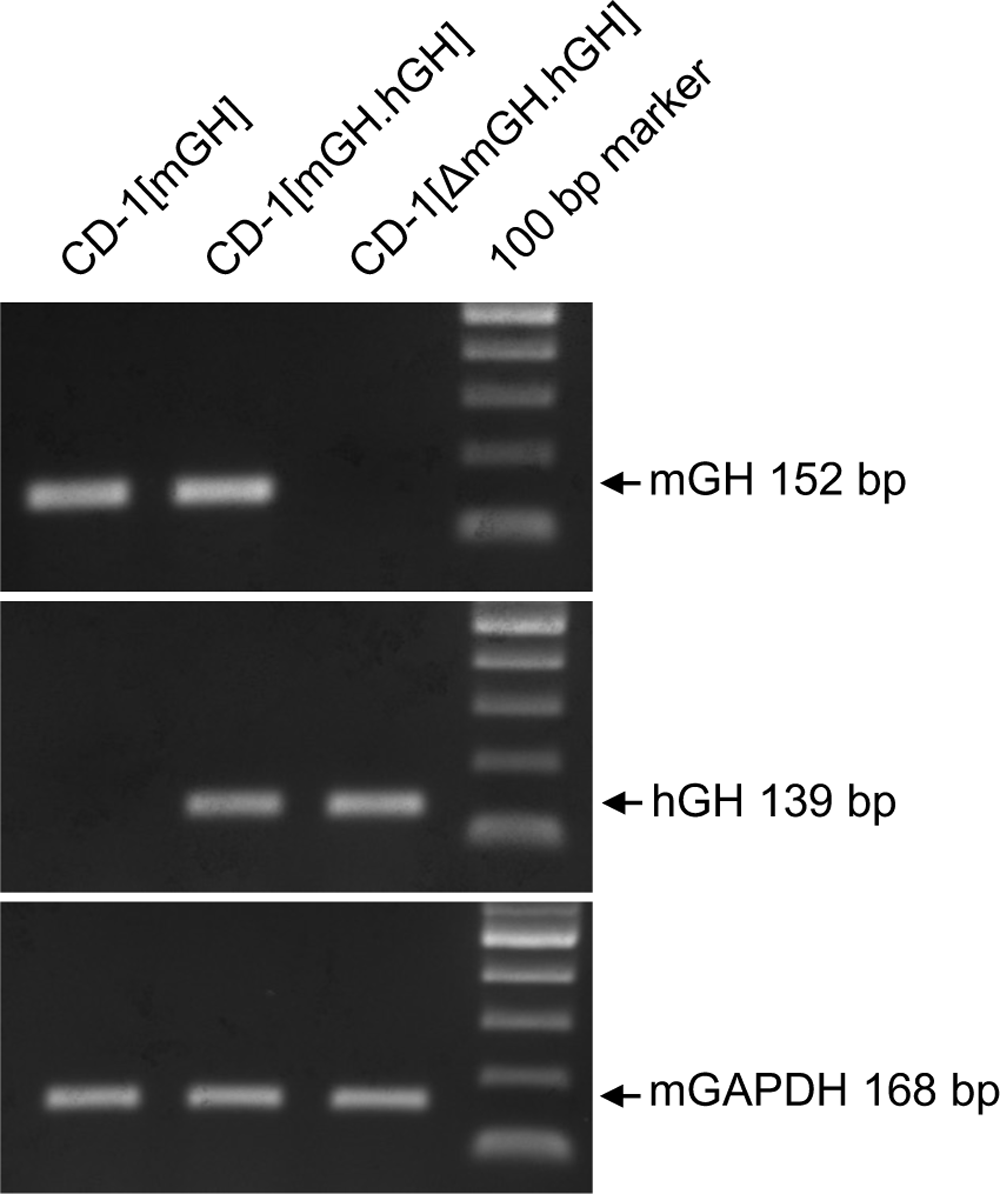
Pituitary growth hormone (GH) gene expression in adult male wild type CD[mGH], transgenic CD-1[mGH.hGH] and CD-1[ΔmGH.hGH] mice. Total RNA was isolated from mouse pituitaries and assessed by PCR using specific primers to amplify mouse (m) growth hormone (GH) and human (h) GH, 1.5% agarose gel electrophoresis and ethidium bromide staining. Mouse glyceraldehyde 3-phosphate dehydrogenase (GAPDH) was also assessed for each sample and used as a control for loading. Size of amplification products are indicated in base pairs (bp).

Circulating hGH levels were assessed in CD-1[ΔmGH.hGH] on RCD and compared to values previously reported for CD-1[mGH.hGH] mice under similar conditions by ELISA [30]. Higher levels of hGH were detected in 8 week-old CD-1[ΔmGH.hGH] mice (0.75 ± 0.21 ng/mL, n=4) *versus* that reported previously for 7 week-old CD-1[mGH.hGH] mice (∼0.05 ng/mL), which are normal in size but also produce ∼12 ng/mL mGH [30]. The level of mGH in 8 week-old wild type CD-1[mGH] mice was 17.61 ± 0.39 ng/ml.

CD-1[ΔmGH.hGH] were visibly smaller (∼20% head to tail) than wild type CD-1[mGH] mice (Figure 2A). In a comparison of mean body weight, CD-1[ΔmGH.hGH] on RCD, weighed ∼42%, ∼30% and ∼31% less than age-matched CD-1[mGH] mice at 4, 12 and 24 weeks, respectively (Figure 2B and C). A similar pattern of increased body weight with time on diet was observed for both CD-1[ΔmGH.hGH] and CD-1[mGH] mice over the 24 weeks on a HFD *versus* RCD. At 4 weeks-old, mean body weight for CD-1[ΔmGH.hGH] mice was 18.1 ± 0.4 g (n=24) but this increased ∼2.6-fold to 47.0 ± 1.1 g after 24 weeks on HFD (n=8). Similarly, at 4 weeks-old, while the mean body weight for CD-1[mGH] mice was 31.3 ± 0.5 g (n=24), a comparable ∼2.1-fold increase to 64.3 ± 2.8 g (n=8) was measured after 24 weeks on HFD.

**Figure 2:**
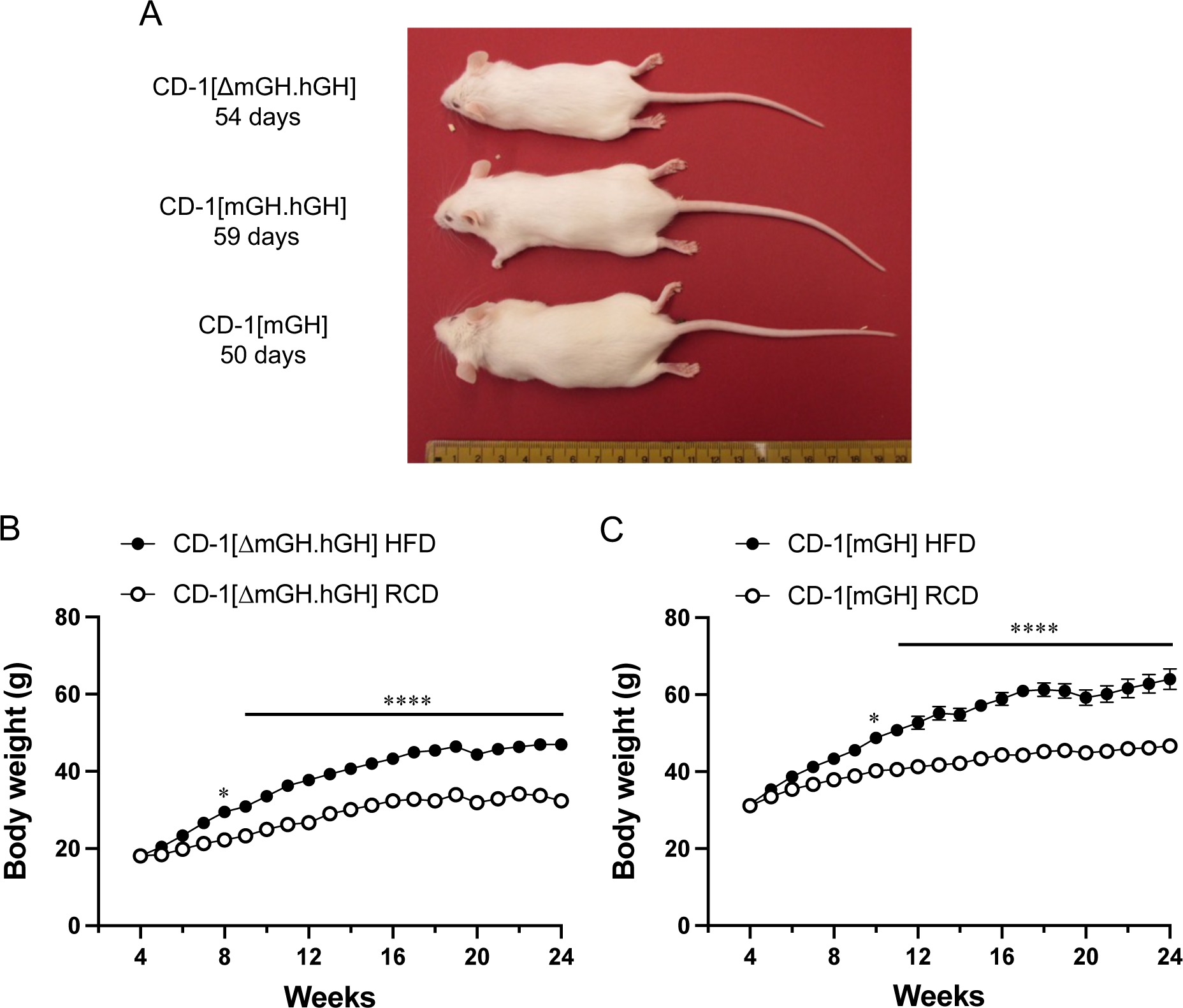
CD-1[ΔmGH.hGH] mouse size and growth on a high fat diet (HFD) or regular chow diet (RCD). (**A**) Male CD-1 mice of similar age (50-59 days) that express human (h) growth hormone (GH) but not mouse (m) GH (CD-1[ΔmGH.hGH] mouse) or both hGH and mGH (CD-1[mGH.hGH] mouse) or mGH (wild type CD-1[mGH] mouse) are shown. The effect of a high fat diet (HFD) or regular chow diet (RCD) on male (**B**) CD-1[ΔmGH.hGH] or (**C**) CD-1[mGH] mouse body weight gain was assessed as a reflection of time in weeks on the diet by two-way ANOVA with Tukey’s *post hoc* test. Values are the mean plus or minus standard error of the mean (*, p<0.05; ****, p<0.0001, n=24).

Considering their smaller size, 8-week-old male CD-1[ΔmGH.hGH] mice on either a HFD or RCD were assessed for their capacity to secrete hGH in response to a GHRP-2 (Figure 3). Four week-old mice were either maintained on RCD or fed a HFD for 4 weeks and their response to a single injection of GHRP-2 was assessed. At 10 minutes post GHRP-2 injection, a significant ∼5.3-fold increase in hGH serum concentration was observed in mice maintained on a RCD (p<0.0001) but did not reach significance in mice fed HFD (p=0.0658) despite an ∼2.8-fold increase (treatment: *F*(1,28)=30.82; *p*<0.0001; diet: *F*(1,28)=2.216; *p*=0.1478; treatment and diet: *F*(1,28)=3.497; *p*=0.072, Figure 3).

**Figure 3.**
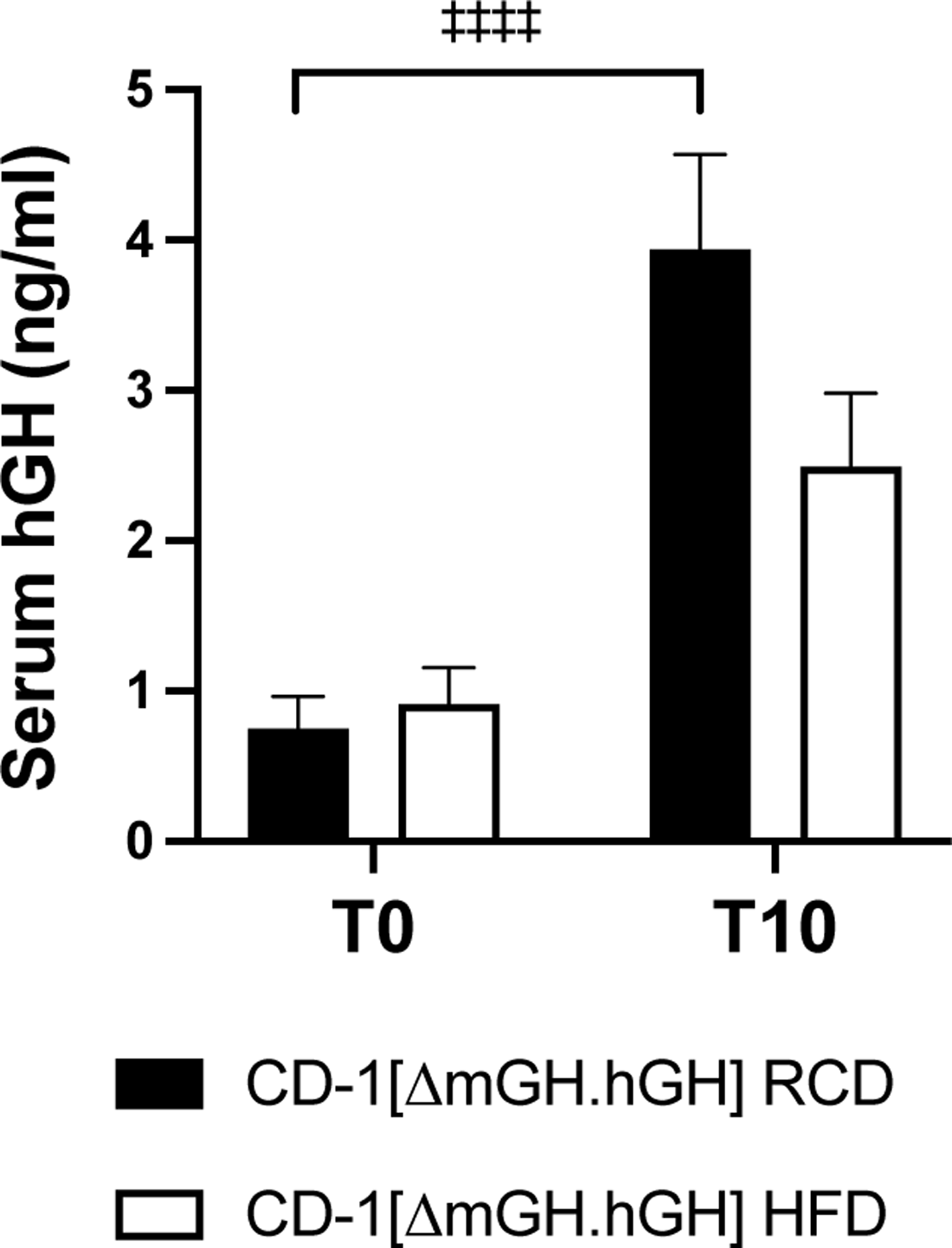
Effect of growth hormone releasing peptide-2 (GHRP-2) on human (h) growth hormone (GH) levels in CD-1[ΔmGH.hGH] mice. Circulating hGH levels were assessed by ELISA in male 8 week-old CD-1[ΔmGH.hGH] mice that had been maintained on either a regular chow diet (RCD) or high fat diet (HFD) for 4 weeks. Serum was sampled at time zero (T0) before the injection with GHRP-2 (0.33 mg/kg) and then 10 minutes post-injection (T10). Results were assessed by two-way ANOVA with Tukey’s *post hoc* test. Values are the mean plus or minus standard error of the mean. A significant difference in hGH levels related to GHRP-2 treatment is indicated by: ‡‡‡‡ p<0.0001 (n=4).

In addition, total RNA was isolated from liver samples of CD-1[ΔmGH.hGH] and wild type CD-1[mGH] mice fed a RCD for 12 and 28 weeks, and assessed for IGF-1 transcripts by qPCR (Figure 4). A similar pattern was seen at both time points. Liver IGF-1 RNA levels were 16% lower at 12 weeks (*p*=0.0113) and 31% lower at 28 weeks (*p*<0.0001) in CD-1[ΔmGH.hGH] mice *versus* wild type CD-1[mGH] mice, respectively, while no effect of HFD (8 and 24 weeks) was observed (mouse type: *F*(1,70)=51.82; *p*<0.0001; time: *F*(1,70)=11.72; *p*=0.001; mouse type and time: *F*(1,70)=5.194; *p*=0.0257, Figure 4).

**Figure 4.**
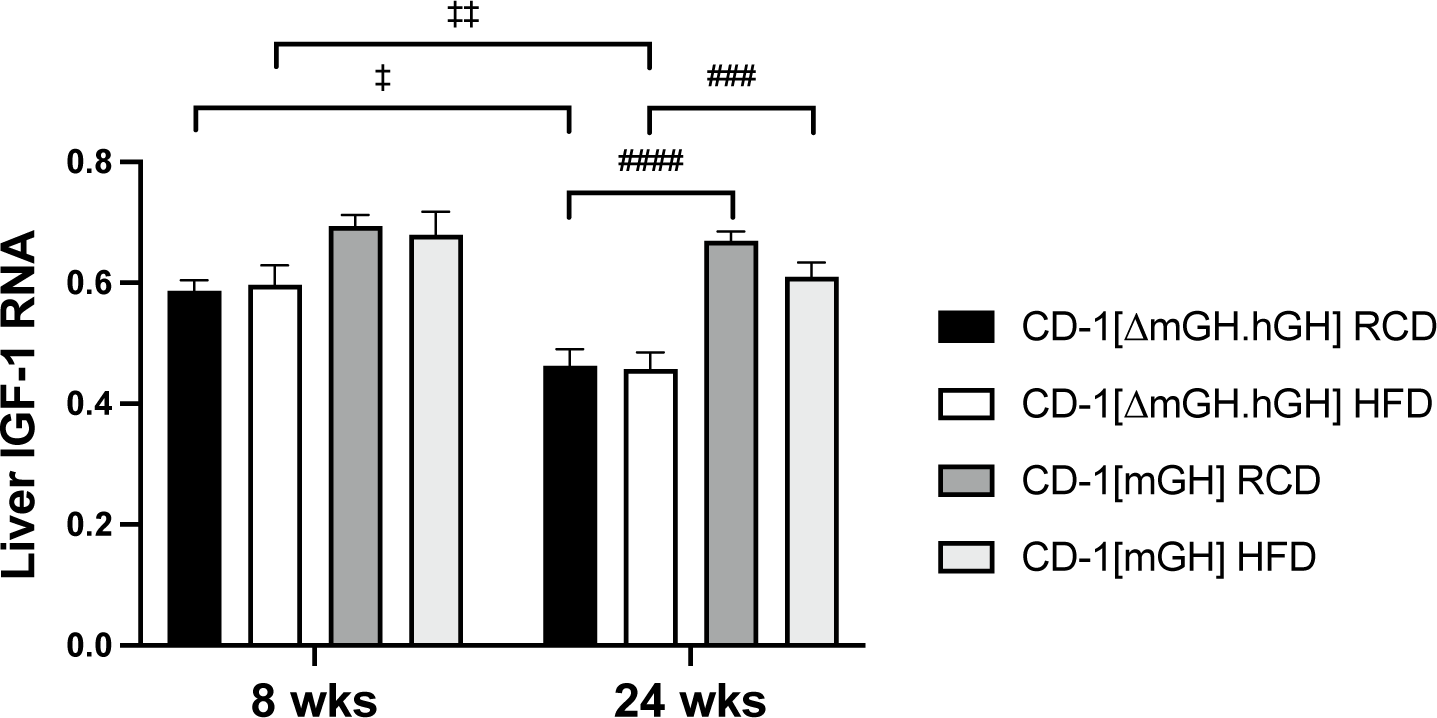
Effect of high fat diet (HFD) compared to a regular chow diet (RCD) on liver IGF-1 RNA levels in male CD-1[ΔmGH.hGH] and wild type CD-1[mGH] mice. Total RNA was isolated from the livers of mice fed HFD for 8 or 24 weeks (wks), or maintained on a RCD. IGF-1 RNA levels relative to GAPDH transcripts were assessed with specific primers by qPCR, and calculated from a standard curve assuming similar reaction kinetics (absolute quantification). Results were assessed by three-way ANOVA with Tukey’s *post hoc* test. Values are the mean plus or minus standard error of the mean. Significant differences related to time (on a diet) are indicated by: ‡ p<0.05; ‡‡ p<0.01 (n=8-12). Significant differences between mouse types are indicated by: ### p<0.001 #### p<0.0001 (n=8-12).

### Effect of HFD on bone density and adiposity in male CD-1[ΔmGH.hGH] mice

Cortical and trabecular bone densities were also assessed in 4 week-old CD-1[ΔmGH.hGH] and CD-1[mGH] mice by micro-CT scanning after 24 weeks on a HFD or maintained on a RCD. There was no significant difference between cortical bone densities in CD-1[ΔmGH.hGH] and wild type CD-1[mGH] mice and also no effect of diet when assessed by two-way ANOVA (cortical bone, diet: *F*(1,25)=0.05708; *p*=0.8131; mouse type: *F*(1,25)=0.002163; *p*=0.9633; diet and mouse type: *F*(1,25)=0.3181; *p*=0.5777; Figure 5A). However, while trabecular bone density was similar in both CD-1[ΔmGH.hGH] and wild type CD-1[mGH] mice, a significant reduction (∼22%, p=0.0271, n=7) in trabecular bone density was seen in CD-1[ΔmGH.hGH] but not CD-1[mGH] mice fed a HFD *versus* RCD (trabecular bone, diet: *F*(1,25)=7.752; *p*=0.0101; mouse type: *F*(1,25)=2.104; *p*=0.1594; diet and mouse type: *F*(1,25)=2.452; *p*=0.13, Figure 5B).

**Figure 5.**
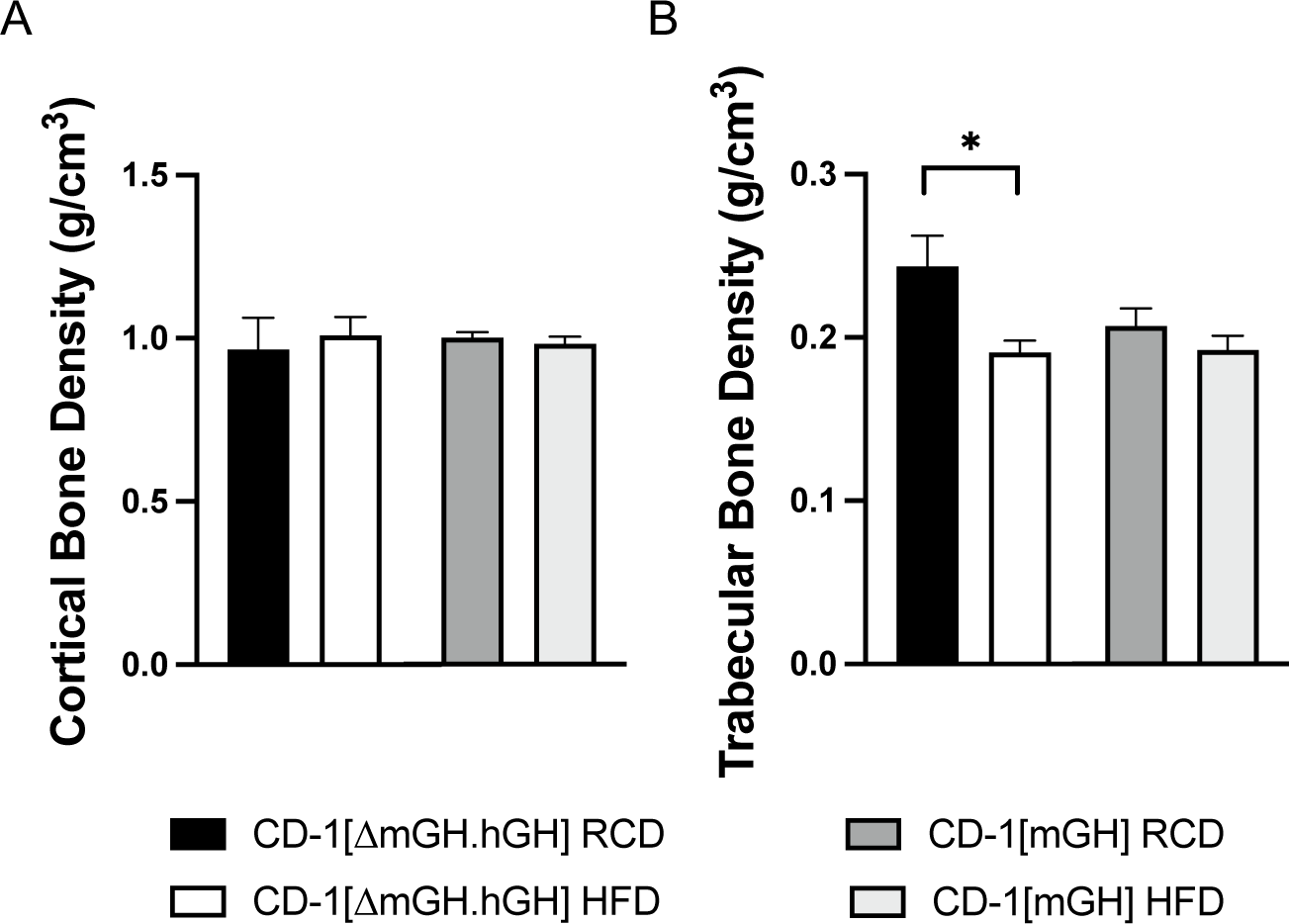
Effect of high fat diet (HFD) compared to a regular chow diet (RCD) on bone density in male CD-1[ΔmGH.hGH] and wild type CD-1[mGH] mice. The densities of cortical and trabecular bone, g (hydroxyapatite)/cm^3^, were measured in male CD-1[ΔmGH.hGH] and wild type CD-1[mGH] mice by micro-CT scanning after 24 weeks on a HFD or maintained on a RCD. Results were assessed by two-way ANOVA with Tukey’s *post hoc* test. Values are the mean plus or minus standard error of the mean. A significant difference related to diet is indicated by: * p<0.05 (n=7).

Visual inspection of body cavities suggested greater abdominal fat in CD-1[ΔmGH.hGH] *versus* CD-1[mGH] mice at 12 weeks of age in mice maintained on RCD (compare panels in Figure 6A and B). A further increase was suggested in CD-1[ΔmGH.hGH] mice fed HFD for 8 weeks, and this diet regimen also resulted in evidence of increased fat deposition in CD-1[mGH] mice (Figure 6B). Total body adiposity was measured by micro-CT in 28 week-old CD-1[ΔmGH.hGH] and CD-1[mGH] mice fed a HFD for 24 weeks or maintained on a RCD. A significant ∼1.7-fold increase in body fat content per body volume was detected in CD-1[ΔmGH.hGH] *versus* CD-1[mGH] mice (*p*=0.0467; Figure 6C). In addition, HFD *versus* RCD resulted in significant increases in total body fat/body volume in CD-1[ΔmGH.hGH] mice (∼1.7-fold increase, *p*=0.0002) and CD-1[mGH] mice (∼2.2-fold increase, *p*=0.0002) (diet: *F*(1,26)=49.59; *p*<0.0001; mouse type: *F*(1,26)=14.95; *p*=0.0007; diet and mouse type: *F*(1,26)=0.003384; *p*=0.9541, Figure 6C).

**Figure 6.**
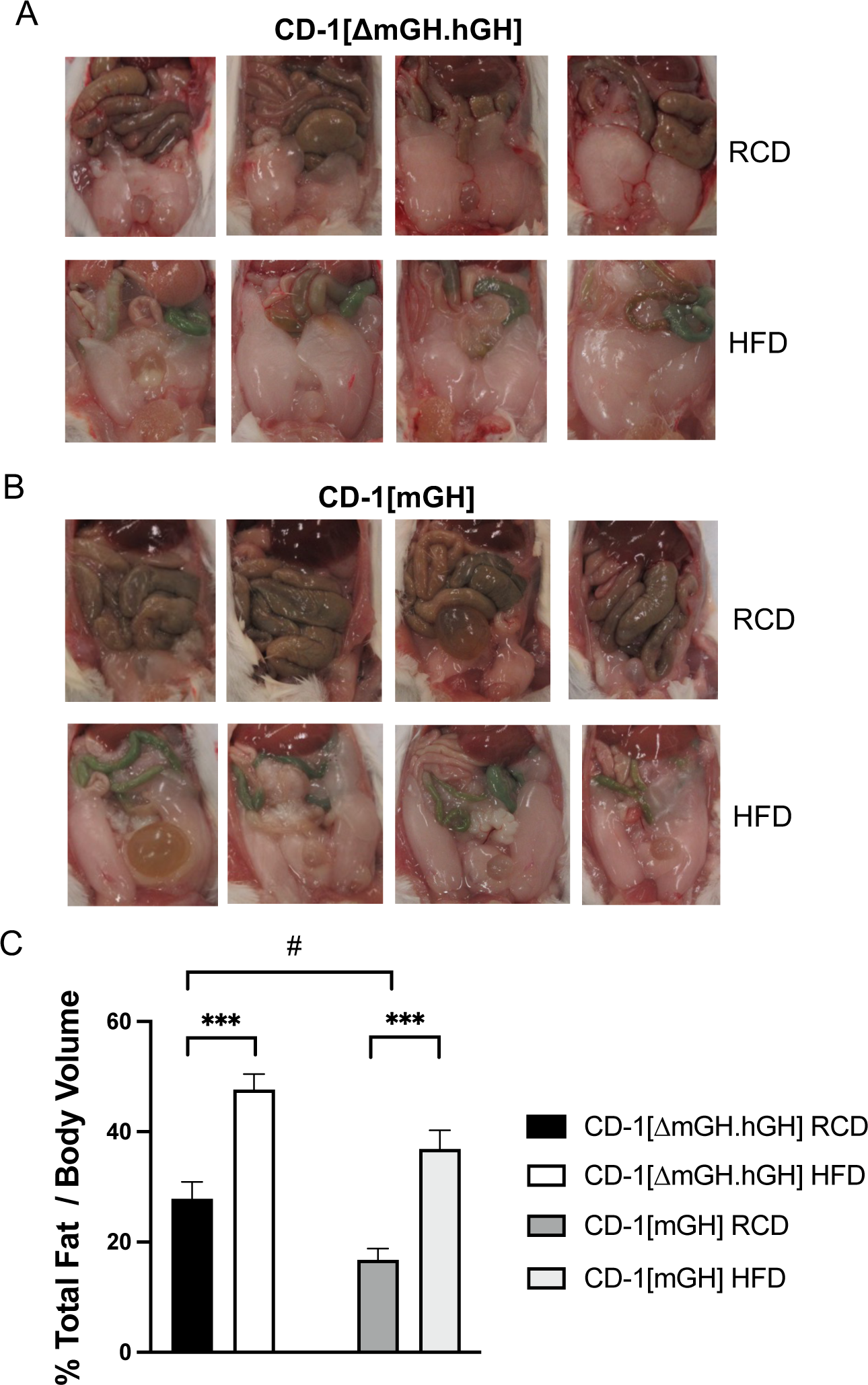
Assessment of body fat. Comparison of abdominal cavities from 12 week-old male (**A**) CD-1[ΔmGH.hGH] and (**B**) wild type CD-1[mGH] mice fed high fat diet (HFD) for 8 weeks or maintained on a regular chow diet (RCD). (**C**) Total body adiposity was measured by micro-CT in 28 week-old male CD-1[ΔmGH.hGH] and CD-1[mGH] mice fed a HFD for 24 weeks or maintained on a RCD. Results were assessed by two-way ANOVA with Tukey’s *post hoc* test. Values are the mean plus or minus standard error of the mean. Significant differences related to diet are indicated by: *** p<0.05 (n=7-8). A significant difference related to mouse type is indicated by: # p<0.05 (n=7-8).

### Effect of HFD on glucose clearance in male CD-1[ΔmGH.hGH] mice

A glucose tolerance test (GTT) was done on 12 week-old CD-1[ΔmGH.hGH] and CD-1[mGH] mice that were fed HFD for 8 weeks as well as 28 week-old mice that were fed HFD for 24 weeks, and compared to mice maintained on RCD. The pattern of glucose clearance was assessed for up to 120 minutes post glucose injection (Figure 7). A HFD for 8 weeks significantly impaired glucose clearance in CD-1[mGH] but not CD-1[ΔmGH.hGH] mice, including the comparison of glucose levels at 120 minutes post injection (CD-1[ΔmGH.hGH] mouse, diet: *F*(1,38)=3.242; *p*=0.0797; time: *F*(6,38)=12.2, *p*<0.0001; diet and time: *F(*6,38)=0.7006, *p*=0.6508; Figure 7A; CD-1[mGH] mouse, diet: *F*(1,40)=36.14; *p*=0.0001; time: *F*(6,40)=44.8, *p*<0.0001; diet and time: *F(*6,40)=4.684, *p*=0.0011; Figure 7B). One of the four CD-1[mGH] mice assessed still had a glucose level at the limit of detection (33.3 mM) at 90 and 120 minutes post glucose injection. By contrast, HFD for 24 weeks was associated with significant impairment in glucose clearance in both CD-1[ΔmGH.hGH] and CD-1[mGH] mice (CD-1[ΔmGH.hGH] mice, diet: *F*(1,84)=39.48; *p*<0.0001; time: *F*(6,84)=37.63, *p*<0.0001; diet and time: *F(*6,84)=5.63, *p*<0.0001, Figure 7C; and CD-1[mGH] mice, diet: *F*(1,86)=24.22; *p*<0.0001; time: *F*(6,86)=89.9, *p*<0.0001; diet and time: *F(*6,86)=5.203, *p*=0.0001, Figure 7D). Again, one of the seven CD-1[mGH] mice assessed in this group had a glucose level the limit of detection at 90 minutes post injection.

**Figure 7.**
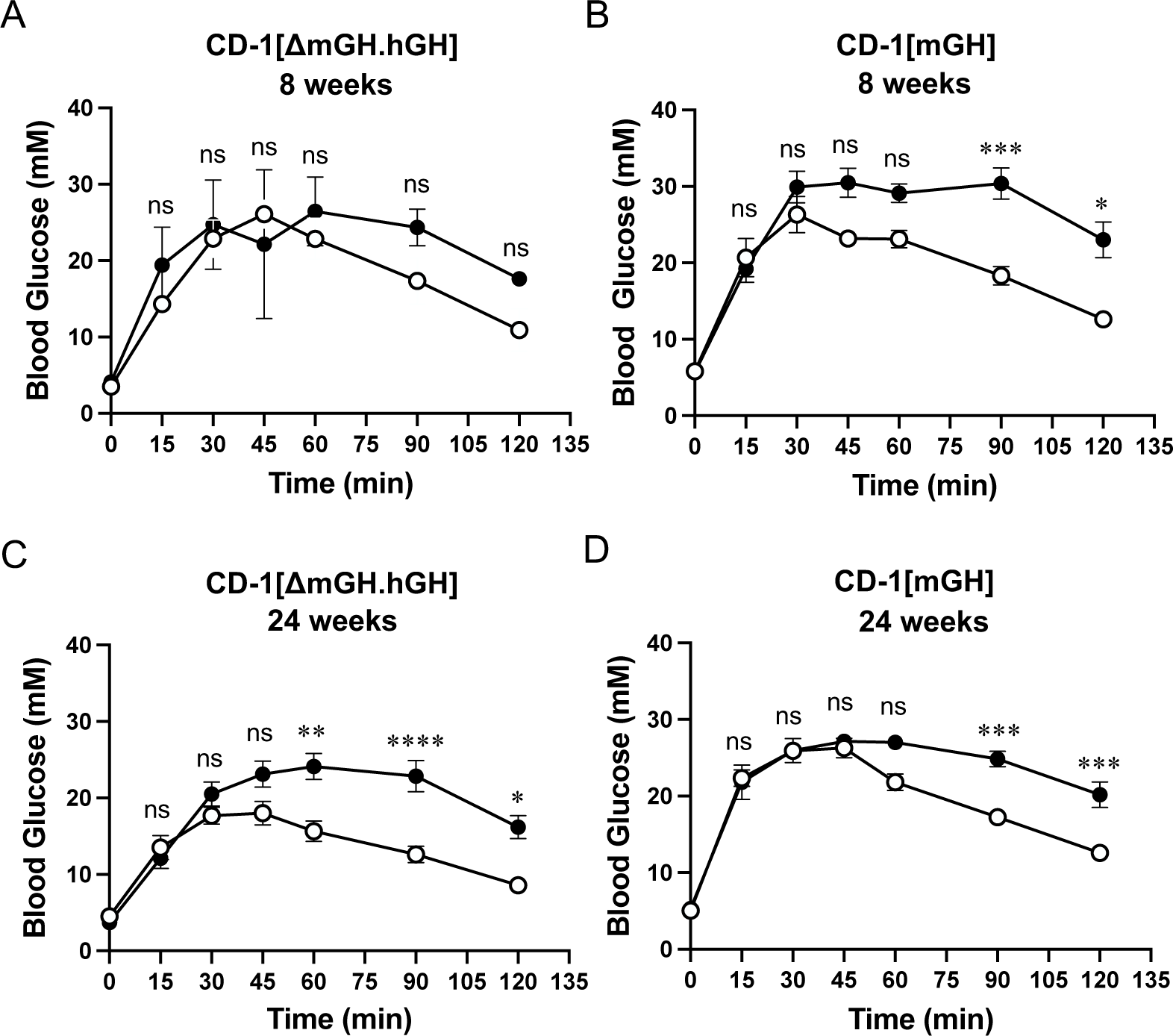
Glucose clearance assessed by glucose tolerance test in (**A** and **C**) CD-1[ΔmGH.hGH] and (**B** and **D**) wild type CD-1[mGH] mice on a high fat diet (HFD; closed circles) or regular chow diet (RCD; open circles). Glucose clearance was assessed over a period of 120 minutes after 8 (**A** and **B**) and 24 (**C** and **D**) weeks on their respective diets. Values are expressed as mean plus or minus standard error of the mean and were determined and analyzed by two-way ANOVA with Tukey’s *post hoc* test. Sample size (n): CD-1[ΔmGH.hGH]/8 weeks=11; CD-1[mGH]/8 weeks=12; CD-1[ΔmGH.hGH]/24 weeks=8; CD-1[mGH]/24 weeks=11. \**p* < 0.05, \*\**p* < 0.01, \*\*\**p* < 0.001; ns, not significant.

### Effect of HFD on pancreatic islet area as well as insulin and IGF-2 gene expression in male CD-1[ΔmGH.hGH] mice

Islet area was assessed in multiple H&E stained sections through the pancreas of CD-1[ΔmGH.hGH] and CD-1[mGH] mice fed a HFD for 8 and 24 weeks or maintained on a RCD. There was no significant effect in 12 week-old mice fed a HFD *versus* RCD for 8 weeks. However, a relative >2-fold increase in islet area was suggested in either 28 week-old CD-1[ΔmGH.hGH] or CD-1[mGH] mice fed a HFD for 24 weeks, however, only the increase in CD-1[ΔmGH.hGH] mice was significant when data from both mouse types were compared (diet: *F*(1,873)=24.45; *p*<0.0001; time: *F*(1,873)=30.31; *p*<0.0001; mouse type: *F*(1,873)=23.63; *p*<0.0001; diet and time: *F*(1,873)=7.07; *p*=0.008; diet and mouse type: *F*(1,873)=1.104; *p*=0.2937; time and mouse type: *F*(1,873)=12.88; *p*=0.0004; diet, time and mouse type: *F*(1,873)=3.033; *p*=0.0819, Figure 8A).

**Figure 8.**
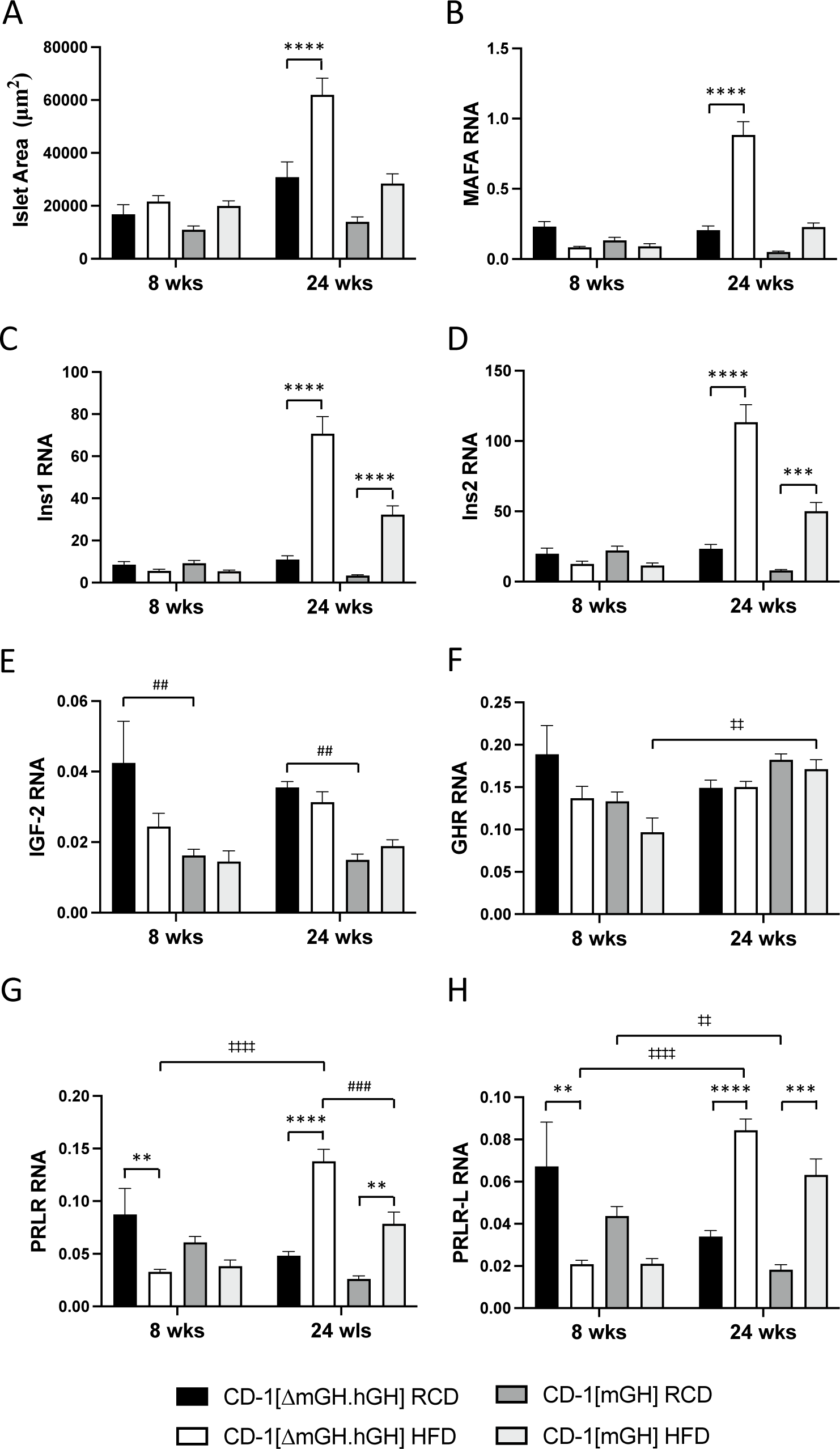
Assessment of pancreatic islet area and RNA levels in male CD-1[ΔmGH.hGH] and wild type CD-1[mGH] mice fed high fat diet (HFD) for 8 or 24 weeks (wks), or maintained on a regular chow diet (RCD). (**A**) Islet area (μm^2^) was determined from digital images of hematoxylin and eosin-stained sections of pancreas using image analysis software. (**B-H**) Total RNA from mouse pancreas was assessed relative to GAPDH transcripts with specific primers by qPCR for: (**B**) MAF bZIP transcription factor A (MAFA), (**C**) insulin/Ins1, (**D**) insulin/Ins2, (**E**) insulin-like growth factor-2 (IGF-2), (**F**) GH receptor (GHR), (**G**) all PRL receptor (PRLR) isoforms and (**H**) the long PRLR isoform (PRLR-L). Results were assessed by three-way ANOVA with Tukey’s *post hoc* test. Values are the mean plus or minus standard error of the mean. Significant differences related to diet are indicated by: ** p<0.01, *** p<0.001, **** p<0.0001 (n=4-8). Significant differences related to time (on the diet) are indicated by: ‡‡ p<0.01, ‡‡‡‡ p<0.0001 (n=4-8). Significant differences related to mouse type are indicated by: ## p<0.01, ### p<0.001 (n=4-8).

Total RNA was isolated from pancreas samples from CD-1[ΔmGH.hGH] and CD-1 [mGH] mice fed a HFD *versus* RCD for 8 weeks and 24 weeks. Transcript levels were determined by qPCR for the: glucose-regulated and pancreatic β-cell-specific transcription factor MAFA and insulin (Ins1 and Ins2) as well as GHR, PRLR and IGF-2 genes. Each RNA data set was analysed by three-way ANOVA to compare between mouse types (CD-1[ΔmGH.hGH] and CD-1[mGH]) as well as the effect of 8 and 24 weeks HFD *versus* RCD in each mouse type. MAFA RNA levels were similar in 12 week-old CD-1[ΔmGH.hGH] and wild type CD-1[mGH] mice fed a HFD *versus* RCD for 8 weeks (Figure 8B). By contrast, an increase in MAFA transcript levels (>4-fold) was suggested in both 28 week-old CD-1[ΔmGH.hGH] or CD-1[mGH] mice fed a HFD for 24 weeks, however, as with islet area, only the increase in CD-1[ΔmGH.hGH] mice was significant when data from both mouse types were compared (MAFA RNA, diet: *F*(1,78)=21.58; *p*<0.0001; time: *F*(1,78)=33.76; *p*<0.0001; mouse type: *F*(1,78)=39.66; *p*<0.0001; diet and time: *F*(1,78)=53.45; *p*<0.0001; diet and mouse type: *F*(1,78)=7.671; *p*=0.007; time and mouse type: *F*(1,78)=25.49; *p*<0.0001; diet, time and mouse type: *F*(1,78)=18.03; *p*<0.0001, Figure 8B).

As with MAFA transcripts, Ins1 and Ins2 RNA levels were similar in both 12 week-old CD-1[ΔmGH.hGH] and wild type CD-1[mGH] mice regardless of diet (Figure 8C and D). However, significant >6-fold increases in both Ins1 and Ins2 transcript levels were seen in both 28 week-old CD-1[ΔmGH.hGH] and wild type CD-1[mGH] mice fed a HFD for 24 weeks *versus* maintenance on RCD, when data from both mouse types were compared (Ins1 RNA, diet: *F*(1,78)=43.13; *p*<0.0001; time: *F*(1,78)=50.84; *p*<0.0001; mouse type: *F*(1,78)=13.33; *p*=0.0005; diet and time: *F*(1,78)=58.84; *p*<0.0001; diet and mouse type: *F*(1,78)=6.493; *p*=0.0128; time and mouse type: *F*(1,78)=13.99; *p*=0.0003; diet, time and mouse type: *F*(1,78)=5.775; *p*=0.0186; and Ins2 RNA, diet: *F*(1,78)=34.75; *p*<0.0001; time: *F*(1,78)=44.17; *p*<0.0001; mouse type: *F*(1,78)=15.99; *p*=0.0001; diet and time: *F*(1,78)=60.08; *p*<0.0001; diet and mouse type: *F*(1,78)=7.054; *p*=0.0096; time and mouse type: *F*(1,78)=17.07; *p*<0.0001; diet, time and mouse type: *F*(1,78)=5.252; *p*=0.0246, Figure 8C and D).

Significantly higher (>2-fold) IGF-2 RNA levels were detected in CD-1[ΔmGH.hGH] when compared to wild type CD-1[mGH] mice maintained on RCD for 12 and 28 weeks (Figure 8E). However, in contrast to MAFA, insulin and PRLR RNA levels, there was no significant effect of diet on IGF-2 transcript levels in either CD-1[ΔmGH.hGH] or wild type CD-1[mGH] mice (IGF-2 RNA, diet: *F*(1,77)=3.124; *p*=0.0811; time: *F*(1,77)=0.0727; *p*=0.7882; mouse type: *F*(1,77)=37.05; *p*<0.0001; diet and time: *F*(1,77)=2.934; *p*=0.0908; diet and mouse type: *F*(1,77)=4.603; *p*=0.0351; time and mouse type: *F*(1,77)=0.07945; *p*=0.7788; diet, time and mouse type: *F*(1,77)=0.5297; *p*=0.4689, Figure 8E).

There was no significant effect of age (12 and 28 week-old) on GHR RNA levels in either CD-1[ΔmGH.hGH] or wild type CD-1[mGH] mice fed RCD, or an effect of HFD in either mouse type (GHR RNA, diet: *F*(1,76)=6.276; *p*=0.0144; time: *F*(1,76)=6.054; *p*=0.0161; mouse type: *F*(1,76)=1.134; *p*=0.2904; diet and time: *F*(1,76)=3.988; *p*=0.0494; diet and mouse type: *F*(1,76)=0.00701; *p*=0.9335; time and mouse type: *F*(1,76)=14.52; *p*=0.0003; diet, time and mouse type: *F*(1,76)=0.4914; *p*=0.4855, Figure 8F). By contrast, effects of HFD on PRLR (long and short isoforms) as well as more specifically transcript levels for the long PRLR isoform (PRLR-L) were observed in both CD-1[ΔmGH.hGH] and wild type CD-1[mGH] mice (Figure 8F and G). However, while a decrease in PRLR RNA levels was suggested in 12 week-old mice after 8 weeks HFD, significant >2-fold increases in RNA levels were detected in 28 week-old mice fed HFD for 24 weeks, which correlate with increased insulin gene expression (PRLR RNA, diet: *F*(1,78)=5.077; *p*=0.027; time: *F*(1,78)=6.058; *p*=0.0161; mouse type: *F*(1,78)=12.62; *p*=0.0007; diet and time: *F*(1,78)=57.55; *p*<0.0001; diet and mouse type: *F*(1,78)=0.03419; *p*=0.8538; time and mouse type: *F*(1,78)=4.33; *p*=0.0407; diet, time and mouse type: *F*(1,78)=5.706; *p*=0.0193, Figure 8G; and PRLR-L RNA, diet: *F*(1,78)=1.637; *p*=0.2046; time: *F*(1,78)=5.187; *p*=0.0255; mouse type: *F*(1,78)=8.516; *p*=0.0046; diet and time: *F*(1,78)=63.39; *p*<0.0001; diet and mouse type: *F*(1,78)=0.7908; *p*=0.3766; time and mouse type: *F*(1,78)=0.4356; *p*=0.5112; diet, time and mouse type: *F*(1,78)=1.981; *p*=0.1632, Figure 8H).

### Effect of HFD on serum insulin and triglyceride levels in male CD-1[ΔmGH.hGH] mice

A determination of insulin levels by ELISA and a lipid profile test, including low density lipoprotein (LDL), high density lipoprotein (HDL), total cholesterol (tCHOL) and triglyceride (TG), were done on serum samples taken from 12 week-old CD-1[ΔmGH.hGH] and wild type CD-1[mGH] mice fed a HFD *versus* RCD for 8 weeks. Each data set was analysed by two-way ANOVA to compare the effect of diet (8 weeks HFD *versus* maintenance on RCD) between mouse types (CD-1[ΔmGH.hGH] and CD-1[mGH]) (Figure 9). There was no significant difference in serum insulin levels in CD-1[ΔmGH.hGH] compared to CD-1[mGH] mice maintained on a RCD. However, >6-fold increases were suggested with HFD *versus* RCD in both mouse types but only reached significance in CD-1[mGH] mice (serum insulin, diet: *F*(1,28)=14.99; *p*=0.0006; mouse type: *F*(1,28)=4.019; *p*=0.0547; diet and mouse type: *F*(1,28)=2.173; *p*=0.1516, Figure 9A).

**Figure 9.**
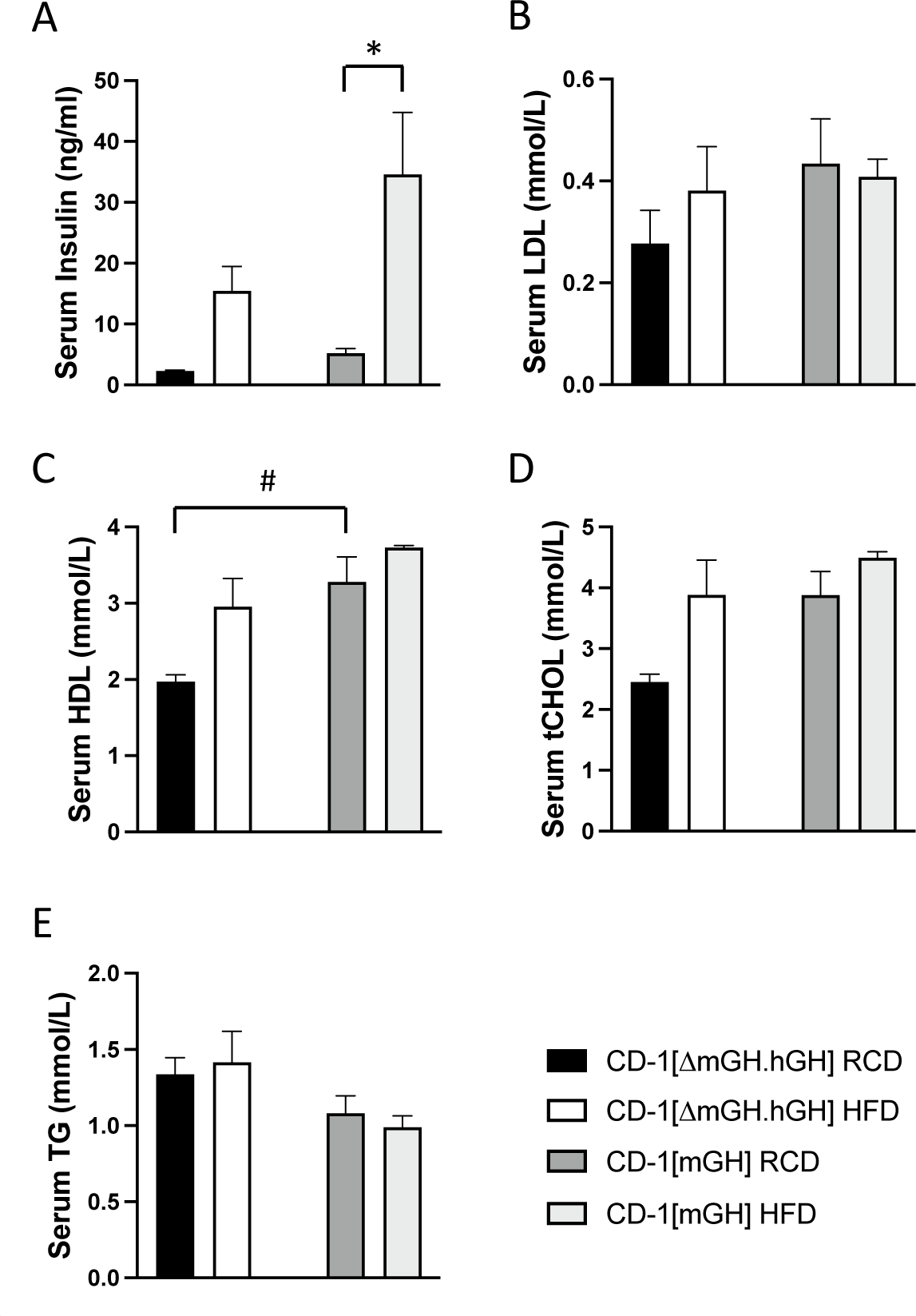
Assessment of serum (**A**) insulin, (**B**) low density lipoprotein-cholesterol (LDL), (**C**) high density lipoprotein-cholesterol (HDL), (**D**) total cholesterol (tCHOL) and (**E**) triglyceride (TG) levels in 12-week-old male CD-1[ΔmGH.hGH] and wild type CD-1[mGH] mice fed high fat diet (HFD) for 8 weeks or maintained on a regular chow diet (RCD). Values are expressed as mean plus or minus standard error of the mean and were determined and analyzed by two-way ANOVA with Tukey’s *post hoc* test. Significant differences related to diet are indicated by: * p<0.05 (n=4-8). Significant differences related to mouse type are indicated by: # p<0.05 (n=4-8).

A similar pattern was observed for LDL, HDL and tCHOL in both mouse types and for the effect of HFD (compare Figures 9B-D). Specifically, lower levels of LDL, HDL and tCHOL were suggested in serum samples from CD-1[ΔmGH.hGH] compared to CD-1[mGH] mice fed RCD for 12 weeks but only reached significance for HDL. In addition, a modest (<2-fold) but not significant increase in levels of LDL, HDL and tCHOL were suggested in CD-1[ΔmGH.hGH] but not CD-1[mGH] fed a HFD for 8 weeks, reaching levels similar to those detected in CD-1[mGH] mice (LDL, diet: *F*(1,12)=0.2967; *p*=0.5959; mouse type: *F*(1,12)=1.666; *p*=0.2211; diet and mouse type: *F*(1,12)=0.8327; *p*=0.3795, Figure 9B; HDL, diet: *F*(1,12)=8.061; *p*=0.0149; mouse type: *F*(1,12)=17.09; *p*=0.0014; diet and mouse type: *F*(1,12)=1.098; *p*=0.3154, Figure 9C; tCHOL, diet: *F*(1,12)=8.387; *p*=0.0134; mouse type: *F*(1,12)=8.325; *p*=0.0137; diet and mouse type: *F*(1,12)=1.365; *p*=0.2654; Figure 9D).

There was no significant effect of diet on serum TG (Figure 9E) in either CD-1[ΔmGH.hGH] or CD-1[mGH] mice. Although higher levels of serum TG were suggested in CD-1[ΔmGH.hGH] compared to CD-1[mGH] mice, this did not reach significance (serum TG, diet: *F*(1,12)=0.0023; *p*=0.9629; mouse type: *F*(1,12)=6.506; *p*=0.0254; diet and mouse type: *F*(1,12)=0.404; *p*=0.537, Figure 9E).

### Estimate of liver discoloration and TG content

Liver discoloration was assessed in 12 and 28 week-old CD-1[ΔmGH.hGH] and CD-1[mGH] mice after 8 and 24 weeks on HFD, respectively. A six point (0-5) scale spanning a healthy dark red liver (score of 1 or less) to an unhealthy pink liver (score of greater than 3) was used in a blind-study to assess digital images of mouse livers and derive a mean score (Figure 10A). No difference in the pattern of liver coloration was observed between CD-1[ΔmGH.hGH] and CD-1[mGH] mice, and both mouse types displayed similar levels of discoloration, consistent with increased fat content and steatosis, after 8 and 24 weeks of HFD *versus* RCD.

**Figure 10.**
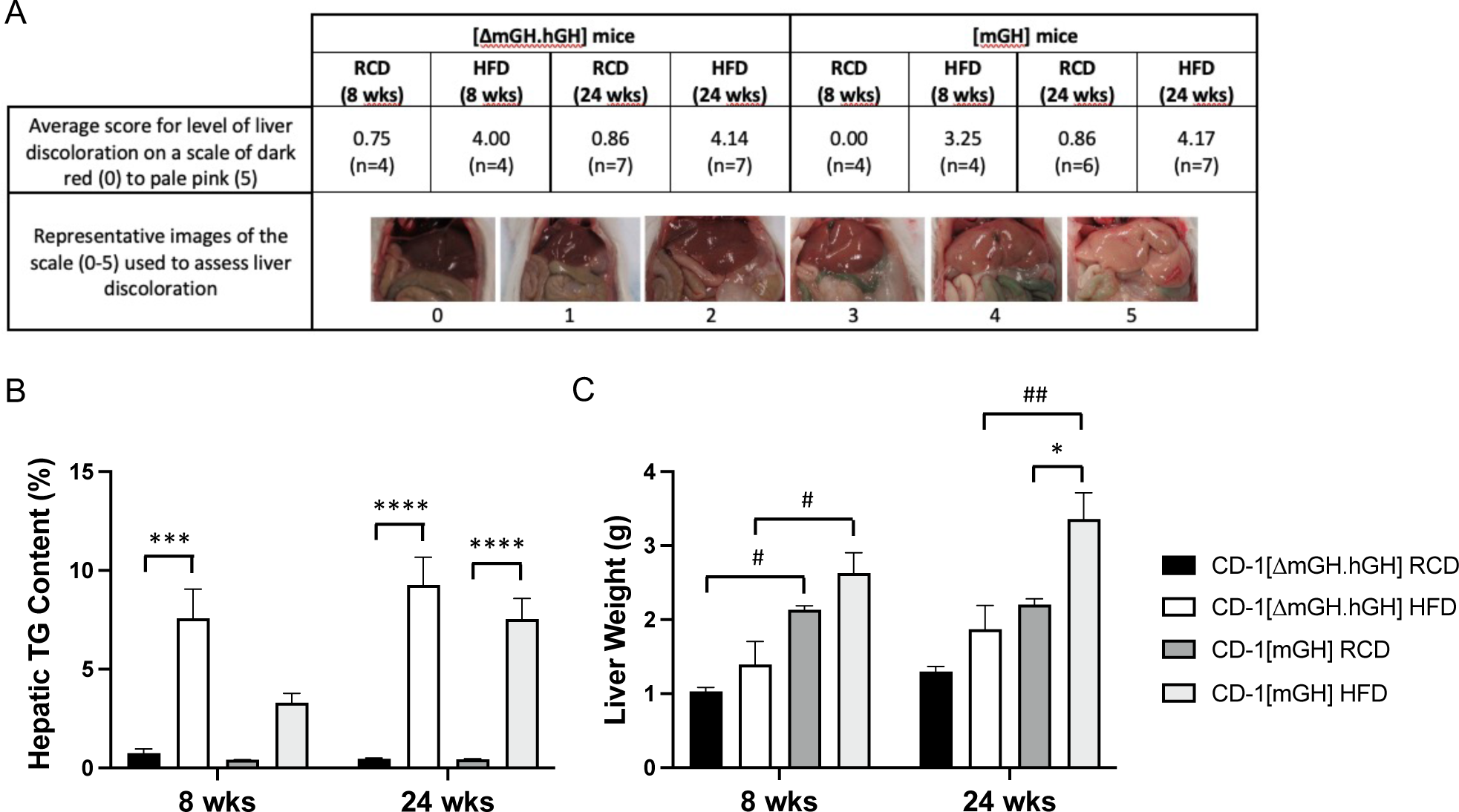
Assessment of fatty liver/steatosis through: (**A**) an estimate of liver discoloration in a blinded assessment of digital images scored with a numerical scale, (**B**) hepatic triglyceride (TG) content as a percentage of liver weight, as well as a determination of (**C**) liver weight (g) in male CD-1[ΔmGH.hGH] and wild type CD-1[mGH] mice fed high fat diet (HFD) for 8 or 24 weeks (wks), or maintained on a regular chow diet (RCD). Data were assessed by three-way ANOVA with Tukey’s *post hoc* test. Values are the mean plus or minus standard error of the mean. Significant differences related to diet are indicated by: * p<0.05, *** p<0.001, **** p<0.0001 (n=4). A significant difference related to mouse type is indicated by: # p<0.05, ## p<0.01 (n=4).

Liver (hepatic) TG content was determined as a percentage of liver weight in both 12 and 28 week-old CD-1[ΔmGH.hGH] and CD-1[mGH] mice after 8 and 24 weeks, respectively, on HFD or maintained on RCD. Each data set was analysed by three-way ANOVA to compare between mouse types (CD-1[ΔmGH.hGH] and CD-1[mGH]) as well as the effect of 8 and 24 weeks HFD *versus* RCD in each mouse type. A >7-fold increase in liver TG content in both mouse types fed a HFD for 8 or 24 was observed (Figure 10B). However, this increase did not reach significance for wild type CD-1[mGH]) mice after only 8 weeks HFD (diet: *F*(1,24)=119.4; *p*<0.0001; time: *F*(1,24)=5.826; *p*=0.0238; mouse type: *F*(1,24)=7.38; *p*=0.012; diet and time: *F*(1,24)=6.946; *p*=0.0145; diet and mouse type: *F*(1,24)=5.764; *p*=0.245; time and mouse type: *F*(1,24)=1.506; *p*=0.2317; diet, time and mouse type: *F*(1,24)=0.9019; *p*=0.3517, Figure 10B). The mean values for liver TG content exceeded 5% for both mouse types after 24 weeks HFD as well as CD-1[ΔmGH.hGH] mice after 8 weeks HFD but not CD-1[mGH] mice (Figure 10B); 5% liver TG content has been used to suggest hepatic steatosis [41]. Consistent with mouse size, the mean liver weight was significantly (52%) lower in CD-1[ΔmGH.hGH] compared to CD-1[mGH] mice maintained on RCD for 12 weeks; liver weight was also 41% lower in CD-1[ΔmGH.hGH] mice on RCD 28 weeks but did not reach significance (diet: *F*(1,24)=16.18; *p*=0.0005; time: *F*(1,24)=5.729; *p*=0.0249; mouse type: *F*(1,24)=54.15; *p*<0.0001; diet and time: *F*(1,24)=1.812; *p*=0.1909; diet and mouse type: *F*(1,24)=1.24; *p*=0.2766; time and mouse type: *F*(1,24)=0.007961; *p*=0.9296; diet, time and mouse type: *F*(1,24)=0.493; *p*=0.4893, Figure 10C). Furthermore, while an increase in liver weight after 24 weeks on HFD was suggested in both mouse types, this only reached significance in CD-1[mGH] mice (Figure 10C).

### Effect of HFD on eWAT adipocyte area in male CD-1[ΔmGH.hGH] mice

Adipocyte area (µm^2^) was determined from hematoxylin and eosin-stained sections of eWAT from CD-1[ΔmGH.hGH] and CD-1[mGH] mice fed a HFD for 8 or 24 weeks and compared to age-matched mice maintained on RCD. An increase in eWAT adipocyte area was already evident in CD-1[ΔmGH.hGH] compared to CD-1[mGH] mice fed RCD (compare Figure 11A and B). More specifically, adipocyte area was ∼2.3-fold (*p*<0.0001) greater in CD-1[ΔmGH.hGH] compared to wild type CD-1[mGH] mice at both 12 and 28 weeks of age (Figure 11C). Increases in adipocyte area were also detected in both mouse types fed HFD for 8 weeks. A further increase was seen in wild type CD-1[mGH] but not CD-1[ΔmGH.hGH] mice fed HFD for 24 weeks compared to RCD. However, it was noted that there was no significant difference in the size of adipocytes from CD-1[ΔmGH.hGH] mice fed either RCD or HFD with adipocytes from CD-1[mGH] mice fed HFD (diet: *F*(1,1269)=142.4; *p*<0.0001; time: *F*(1,1269)=17.7; *p*<0.0001; mouse type: *F*(1,1269)=156.0; *p*<0.0001; diet and time: *F*(1,1269)=11.22; *p*=0.0008; diet and mouse type: *F*(1,1269)=71.06; *p*<0.0001; time and mouse type: *F*(1,1269)=0.9836; *p*=0.3215; diet, time and mouse type: *F*(1,1269)=0.6078; *p*=0.4357, Figure 11C).

**Figure 11.**
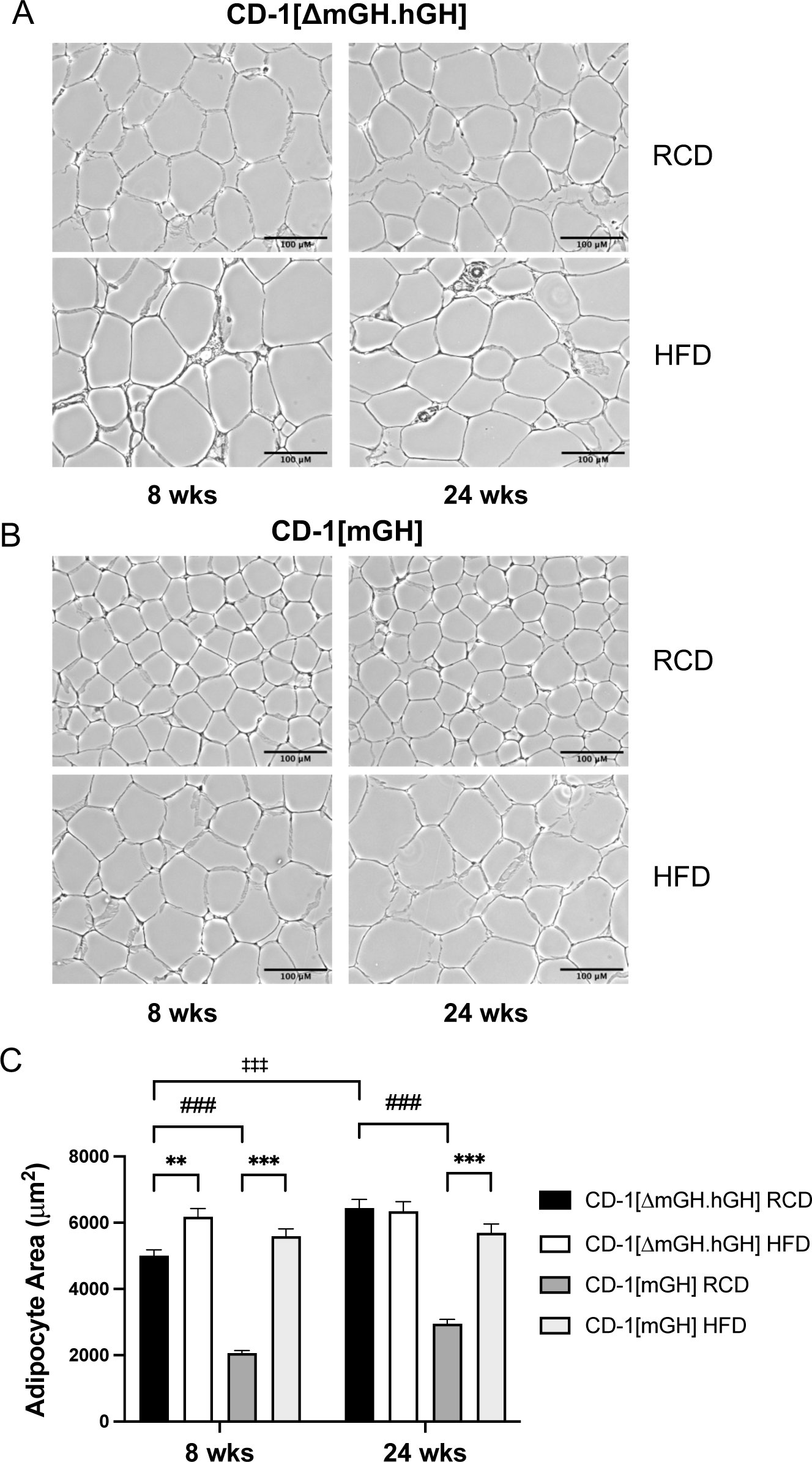
Assessment of epididymal white adipose tissue (eWAT) and adipocyte area. Hematoxylin and eosin-stained sections of eWAT from (**A**) CD-1[ΔmGH.hGH] and (**B**) wild type CD-1[mGH] mice fed high fat diet (HFD) for 8 or 24 weeks (wks), or maintained on a regular chow diet (RCD). (**C**) Adipocyte area (μm^2^) of 40 adipocytes per mouse was captured from digital images of eWAT sections using image analysis software. Data were assessed by three-way ANOVA with Tukey’s *post hoc* test. Values are the mean plus or minus standard error of the mean. Significant differences related to diet are indicated by: ** p<0.01, *** p<0.001 (n=4). A significant difference related to time (on the diet) is indicated by: ‡‡‡ p<0.001 (n=4). A significant difference related to mouse type is indicated by: ### p<0.001 (n=4).

### Effect of HFD on PPARγ, leptin, adiponectin, p16^INK4a^ and p21^CIP1^ eWAT RNA levels in male CD-1[ΔmGH.hGH] mice

Total RNA was isolated from eWAT samples from CD-1[ΔmGH.hGH] and CD-1 [mGH] mice maintained on a HFD *versus* RCD for 8 weeks and 24 weeks. Transcript levels were determined by qPCR for lipolysis-related transcription factor PPARγ, adipokines leptin and adiponectin and senescence-related p16^INK4a^ and p21^CIP1^. Each eWAT RNA data set was analysed by three-way ANOVA to compare between mouse types (CD-1[ΔmGH.hGH] and CD-1[mGH]) as well as the effect of 8 and 24 weeks HFD *versus* RCD in each mouse type (Figure 12). Transcription factor PPARγ RNA levels were similar in both CD-1[ΔmGH.hGH] and wild type CD-1[mGH] mice after 12 and 28 weeks on RCD (Figure 12A). However, significant ∼38% and 42% decreases in PPARγ transcript levels were seen in CD-1[ΔmGH.hGH] (p=0.046) and CD-1[mGH] (*p=*0.0031), respectively, after 24 weeks on a HFD (PPARγ RNA, diet: *F*(1,112)=10.38; *p*=0.0017; time: *F*(1,112)=0.5323; *p*=0.4672; mouse type: *F*(1,112)=3.382; *p*=0.0686; diet and time: *F*(1,112)=16.17; *p*=0.0001, diet and mouse type: *F*(1,112)=0.5221; *p*=0.4714; time and mouse type: *F*(1,112)=0.04757; *p*=0.8277; diet, time and mouse type: *F*(1,112)=3.042; *p*=0.0839, Figure 12A).

**Figure 12.**
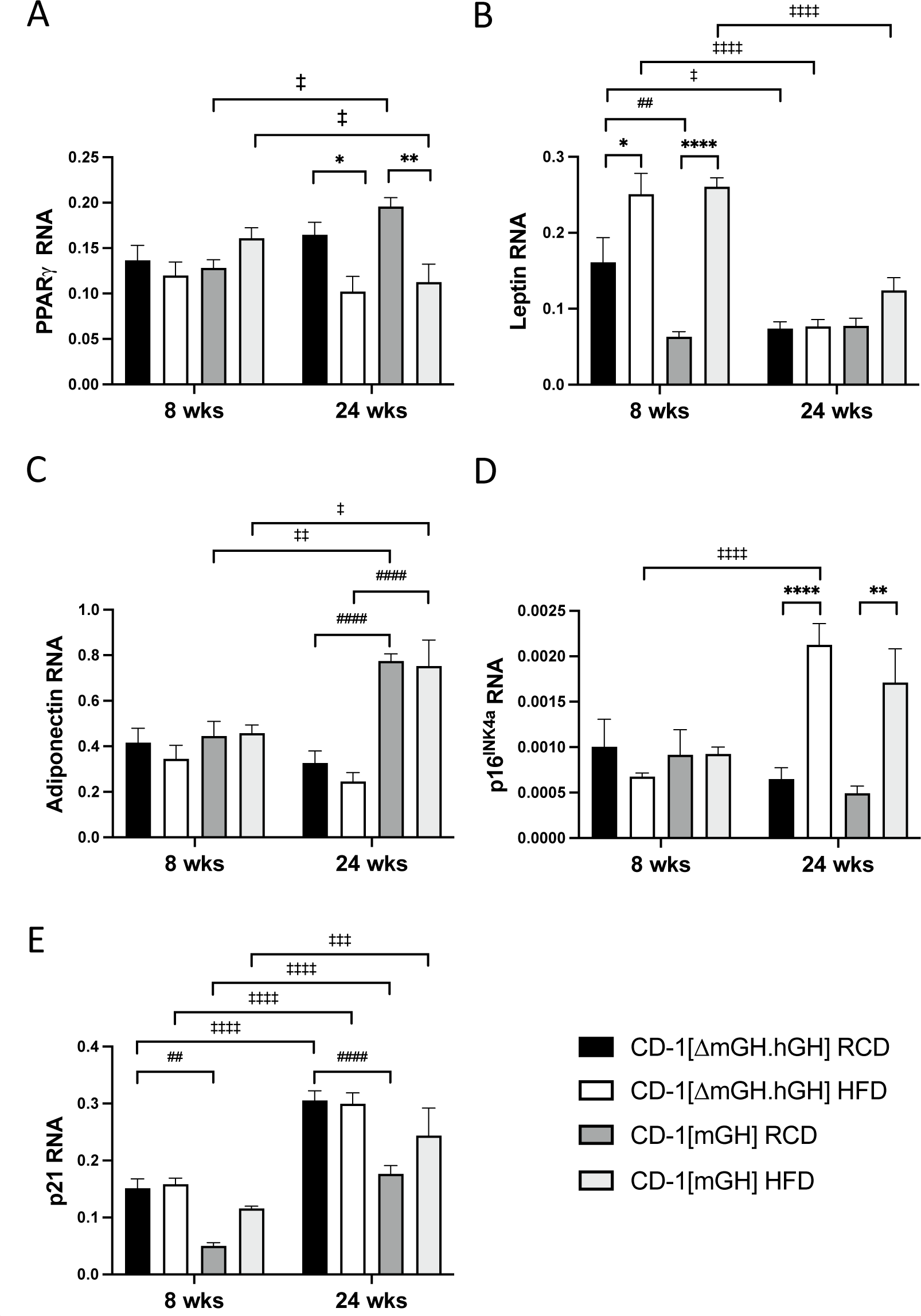
Effect of high fat diet (HFD) compared to a regular chow diet (RCD) on relative epididymal white adipose tissue (WAT) levels of (**A**) peroxisome proliferator-activated receptor γ (PPARγ), (**B**) leptin, (**C**) adiponectin, (**D**) p16^INK4a^ /cyclin-dependent kinase inhibitor 2A and (**E**) p21^CIP1^ RNA from male CD-1[ΔmGH.hGH] and wild type CD-1[mGH] mice. Total RNA from epididymal WAT (eWAT) was isolated from mice maintained on a HFD *versus* RCD for 8 and 24 weeks (wks). Mouse eWAT RNA levels were assessed relative to GAPDH transcripts with specific primers by qPCR. Results were assessed by three-way ANOVA with Tukey’s *post hoc* test. Values are the mean plus or minus standard error of the mean. Significant differences related to diet are indicated by: * p<0.05, ** p<0.01 (n=5-8). Significant differences related to time (on the diet) are indicated by: ‡ p<0.05, ‡‡ p<0.01, ‡‡‡‡ p<0.0001 (n=5-8). Significant differences related to mouse type are indicated by: # p<0.05, ## p<0.01, #### p<0.0001 (n=5-8).

Leptin transcript levels were significantly higher in CD-1[ΔmGH.hGH] compared to wild type CD-1[mGH] mice at 12 weeks (∼2.6-fold, *p=*0.0068) but not 28 weeks on RCD (Figure 12B). In addition, significant increases in leptin RNA levels were detected in wild type CD-1[mGH] mice (∼4.1-fold, p<0.0001) as well as CD-1[ΔmGH.hGH] mice (1.6-fold, *p*=0.0126) after 8 but not 24 weeks on HFD (Leptin RNA, diet: *F*(1,114)=42.07; *p*<0.0001; time: *F*(1,114)=54.68; *p*<0.0001; mouse type: *F*(1,114)=0.5154; *p*=0.4743; diet and time: *F*(1,114)=21.03; *p*<0.0001; diet and mouse type: *F*(1,114)=8.508; *p*=0.0043; time and mouse type: *F*(1,114)=7.167; *p*=0.0085; diet, time and mouse type: *F*(1,114)=1.523; *p*=0.2197, Figure 12B). Adiponectin RNA levels were significantly lower in CD-1[ΔmGH.hGH] compared to wild type CD-1[mGH] mice at 28 (∼58%, *p*<0.0001) but not 12 weeks (Figure 12C). No effect of diet on adiponectin transcript levels was seen in either mouse types (Adiponectin RNA, diet: *F*(1,105)=0.9765; *p*=0.3523; time: *F*(1,105)=7.072; *p*=0.0091; mouse type: *F*(1,105)=44.83; *p*<0.0001; diet and time: *F*(1,105)=0.07479; *p*=0.785; diet and mouse type: *F*(1,105)=0.7736; *p*=0.3811; time and mouse type: *F*(1,105)=24.59; *p*<0.0001, diet, time and mouse type: *F*(1,105)=0.02165; *p*=0.8833, Figure 12C).

Levels of eWAT transcript levels for p16^Ink4a^ and p21^CIP1^ were assessed as markers of senescence-related pathways [9]. Levels of p16^Ink4a^ RNA were similar in CD-1[ΔmGH.hGH] and wild type CD-1[mGH] mice on RCD at both 12 and 28 weeks (Figure 12D). In addition, similar increases in p16^Ink4a^ transcripts were detected in CD-1[ΔmGH.hGH] (2.1-fold, *p*<0.0001) and wild type CD-1[mGH] (3.4-fold, *p*=0.0057) mice, after 24 but not 8 weeks on HFD (p16^Ink4a^ RNA, diet: *F*(1,106)=16.45; *p*<0.0001; time: *F*(1,106)=6.151; *p*=0.0147; mouse type: *F*(1,106)=0.4862; *p*=0.4872; diet and time: *F*(1,106)=26.47; *p*<0.0001, diet and mouse type: *F*(1,106)=0.01855; *p*=0.8919; time and mouse type: *F*(1,106)=1.549; *p*=0.2160, diet, time and mouse type: *F*(1,106)=1.043; *p*=0.3094, Figure 12D).

For p21^CIP1^, transcript levels were higher in CD-1[ΔmGH.hGH] mice when compared to wild type CD-1[mGH] mice on RCD at both 12 (∼3-fold, *p*=0.0019) and 28 weeks (∼1.7-fold, *p*<0.0001) (Figure 12E). Levels were also increased at 28 weeks relative to 12 weeks in both mouse types. However, there was no effect of diet on p21^CIP1^ RNA levels in both mouse types (p21^CIP1^RNA, diet: *F*(1,110)=7.18; *p*=0.0085; time: *F*(1,110)=120.0; *p*<0.0001; mouse type: *F*(1,110)=42.92; *p*<0.0001; diet and mouse type: *F*(1,110)=6.876; *p*=0.01; time and mouse type: F(1,110)=0.6771; p=0.4124; diet, time and mouse type: F(1,110)=0.08088; p=0.7766, Figure 12E).

## DISCUSSION

The transgenic CD-1[mGH.hGH] mouse line meets its requirements for GH through a combination of pituitary mGH and hGH gene expression [27]. Somatotrophs from these mice express either mGH or hGH, or both mGH and hGH [29]. Nonetheless, mGH production is much greater than hGH in these mice [30]. In the current study, we removed endogenous mGH from CD-1[mGH.hGH] mice through breeding with a null mGH (C57BL/6[Gh^tm1Vlcg^]) mouse [34] to generate CD-1[ΔmGH.hGH] mice. These mice are viable and, as expected, continue to express pituitary GH at relatively low levels compared to wild type CD-1[mGH] mice [30]; however, levels of hGH in CD-1[ΔmGH.hGH] are 15-fold greater than detected in CD-1[mGH.hGH]) mice. This is consistent with an increased capacity for hGH production in the absence of mGH, presumably through an ability to respond, at least in part, to feedback via the hypothalamic-pituitary-somatotropic axis. In this context, CD-1[ΔmGH.hGH] mice responded positively to GHRP-2 resulting in (5.3-fold) increased circulating hGH levels. However, the lower basal levels are associated with a 30-45% smaller mouse compared to wild type CD-1[mGH] mouse body weight assessments at 4 and 24 weeks of age. Despite their smaller size, CD-1[ΔmGH.hGH] mice had more body fat and eWAT adipocyte area was significantly larger in CD-1[ΔmGH.hGH] *versus* wild type CD-1[mGH] mice maintained on a RCD. While GH release is known to correlate inversely with intra-abdominal visceral adiposity, decreased GH can further exacerbate visceral fat accumulation because of decreased hormone sensitive lipolysis [42]. By contrast, the decreased circulating levels of hGH had no significant effect on either cortical or trabecular bone density. This may reflect the absence of an early effect of decreased hGH availability on IGF-1 production; specifically, lower liver IGF-1 transcript levels were detected in CD-1[ΔmGH.hGH] mice at 28 weeks but not 12 weeks of age when compared to age-matched wild type CD-1[mGH] mice. Thus, given hGH production, a positive response to GHRP-2, small size, a negative effect on liver IGF-1 RNA levels, increased fat deposition and adipocyte area, CD-1[ΔmGH.hGH] mice maintained on a RCD possess some characteristics of GH-deficiency (GHD).

A comparison of the CD-1[ΔmGH.hGH] mouse with existing examples of GHD, including the growth hormone receptor antagonist (GHA) mouse [43] and GH knockout (GH^−/−^) mouse [34], reveals some similarities. GHA mice express a bovine (b) GH transgene in which glycine at position 119 is replaced with arginine that produces a protein that antagonizes endogenous mGH resulting in decreased GH activity; this has been described as similar to human congenital GHD [9]. These mice are intermediate in size between a wild type and a GH receptor null mouse [7]. The GH^−/−^ mouse, used here to generate the CD-1[ΔmGH.hGH] mouse, produces no GH [34]. Like the CD-1[ΔmGH.hGH] mouse, both GHA and GH^−/−^ mice are characterized by evidence of decreased IGF-1 production, reduced size and weight with increased total fat deposition [44]. An increase in liver discoloration and TG content (steatosis) was suggested in both CD-1[ΔmGH.hGH] and CD-1[mGH] mice fed a HFD for 8 and 24 weeks, which would be consistent with a relative increase in insulin resistance relative to mice maintained on a RCD. The increase in percentage liver TGs was independent of the lower liver weight measured in the smaller CD-1[ΔmGH.hGH] mouse. However, in spite of this, the CD-1[ΔmGH.hGH] showed an initial resistance to the impaired glucose clearance seen in CD-1[mGH] mice. Furthermore, this lack of significant impairment correlated with higher basal pancreatic IGF-2 transcript levels in 12-week-old CD-1[ΔmGH.hGH] mice, while insulin RNA levels were similar in both mouse types. Insulin stimulates insulin gene expression via the insulin receptor (IR)-A in pancreatic islets and IGF-2 has a high affinity for the IR-A receptor [19]. IGF-2 has been described as a metabolic regulator [19]. Evidence based on β-cell-specific knock-out of IGF-2, suggests that a modest increase in IGF-2 could play a positive role in maintaining glucose homeostasis as insulin-resistance develops [19, 20]. However, it is also reported that overexpression of IGF-2 in β-cells can result in pancreatic dysfunction including endoplasmic reticulum stress [19].

Both CD-1[ΔmGH.hGH] and CD-1[mGH] mice displayed impaired glucose clearance after 24 weeks on HFD suggesting insulin resistance has increased and pancreatic function is more compromised, and beyond any proposed initial benefit related to hGH *versus* mGH production. In support, based on MAFA, Ins1 and Ins2 RNA levels, insulin gene expression was increased significantly in both mouse types in response to 24 weeks HFD and reflected in elevated serum insulin levels. There was a correlation of these effects with an increase in PRLR RNA levels, and certainly for the long isoform, but not GHR. Release of GH in to the circulation promotes lipolysis in adipose tissue, leading to enhanced lipid assimilation by the liver and accelerated hepatic TG production. Moreover, it lowers total and LDL cholesterol by inducing LDL clearance [45]. While the level of HDL was significantly decreased in CD-1[ΔmGH.hGH] mice on RCD, all three lipid parameters (LDL, HDL and tCHOL) were increased significantly when these mice were fed HFD that recapitulates GHD. Additionally, liver-specific deletion of the GHR gene in mice resulted in hepatic steatosis not only due to increased lipogenesis but also to decreased TG secretion from the liver [46]. Although serum triglycerides were not increased significantly in response to 24 weeks HFD and lipolysis-related eWAT PPARγ RNA levels were significantly reduced, elevated liver triglycerides as well as evidence of steatosis were detected in both CD-1[ΔmGH.hGH] and CD-1[mGH] mice. Thus, these observations indicate that insulin resistance has increased in both mouse types, and beyond any initial potential benefit due to hGH *versus* mGH production in CD-1[ΔmGH.hGH] mice related to glucose clearance.

It is unlikely that the higher relative levels of pancreatic IGF-2 RNA in hGH-expressing CD-1[ΔmGH.hGH] compared to mGH-expressing CD-1[mGH] mice are due to basal GH levels/activity. CD-1[ΔmGH.hGH] mice are smaller than CD-1[mGH] mice and GH levels are estimated to be ∼95% lower than determined for CD-1[mGH] mice [30]. An alternative possibility is that the increase in IGF-2 transcript levels reflects the ability of hGH, unlike mGH, to activate the PRLR including potentially during the perinatal and/or pubertal periods. There is a positive effect of PRLR receptor signaling on pancreatic development in the early postnatal period in the mouse and IGF-2 is a known target of PRL [18]. Unlike PRL [47], pituitary GH levels are expected to increase postnatally beyond the perinatal period and reach their highest levels in puberty to be followed by an age-related decline in secretion (somatopause) [48]. Thus, it is possible that during postnatal development, including the normal insulin resistance associated with puberty, that hGH through its lactogenic activity has a significant effect on IGF-2 gene expression in CD-1[ΔmGH.hGH] mice. Specifically, elevated IGF-2 gene expression in the pancreas has the potential to influence β-cell function, which may include more capacity to maintain glucose homeostasis in response to developing HFD-induced insulin-resistance. Certainly, puberty has been shown to represent a time during postnatal development where changes in signaling that might improve or protect β-cell function from a risk of even greater or pathological insulin resistance associated with obesity [49].

As stated, adipocyte area was significantly larger in eWAT from CD-1[ΔmGH.hGH] compared to CD-1[mGH] mice maintained on RCD. In addition, there is evidence of adipocyte dysfunction reflected by increased leptin as well as decreased adiponectin RNA levels in eWAT [22–25]. Leptin RNA increased in both CD-1[ΔmGH.hGH] and wild type CD-1[mGH] mice on HFD for 8 weeks, whereas it was unchanged among groups at 24 weeks when maximum adipocyte size was attained. On the other hand, adiponectin RNA was unchanged by model or diet at 8 weeks, yet adiponectin RNA levels doubled in eWAT of the wild type CD-1[mGH] mice at 24 weeks regardless of RCD or HFD. The increase in p16^INK4a^ transcript levels in eWAT also raises the possibility that a HFD for 24 weeks is associated with an increased potential for cell senescence in both CD-1[ΔmGH.hGH] and wild type CD-1[mGH] mice. This contrasts with p21^CIP1^ levels, which appear intrinsically higher in CD-1[ΔmGH.hGH] compared to wild type CD-1[mGH] mice, when measured at 12 and 28 weeks of age on RCD. These higher levels of p21^CIP1^ RNA in CD-1[ΔmGH.hGH] mice are not consistent with a positive association of GH action with accumulation of senescent cells in adipose tissue, given the significantly lower GH levels [9]. However, these higher p21^CIP1^ transcript levels together with the increase p16^INK4a^ RNA levels in mice on a HFD, do support a correlation with obesity, including fat and adipocyte area, with increased cell senescence [9, 50].

Unlike mGH, hGH is able to bind and activate the PRLR and thus the potential exists for lactogenic activity to affect the phenotype of CD-1[ΔmGH.hGH] compared to CD-1[mGH] mice [10]. In addition, PRLRs have been detected in mouse adipose tissue [51, 52]. Our observations are limited to assessments of size, bone density, body fat, eWAT as well as liver discoloration, glucose clearance and the pancreas in male CD-1[ΔmGH.hGH] mice. The effects observed, with the exception of glucose clearance and the pancreas are consistent with GH insufficiency and thus largely predicted based on the reported phenotype of GHA and GH^−/−^ mice. Similarly, CD-1[ΔmGH.hGH] and GHA mice share decreased reproductive capacity. While this suggests the reduced litter size seen in CD-1[ΔmGH.hGH] mice is not related to any lactogenic activity associated with hGH *versus* its signaling via the GHR, a contribution related to additional lactogenic activity cannot be excluded. The initial resistance to the negative effects of HFD on glucose clearance in hGH-expressing CD-1[ΔmGH.hGH] *versus* wild type CD-1[mGH] mice and specifically a possible link to elevated IGF-2 production, raise the possibility of a differential effect of hGH *versus* mGH during pancreas development. Furthermore, the correlation between PRLR and not GHR with elevated MAFA, Ins1 and Ins2 RNA levels at 24 weeks HFD also support a role for lactogenic signaling in the pancreas to which hGH but not mGH may contribute.

In summary, male CD-1[ΔmGH.hGH] mice produce a lower basal level of GH than wild type CD-1[mGH] mice but possess a responsive hypothalamic-pituitary axis. While these mice are smaller and show signs of GH insufficiency relative to their wild type counterparts, anabolic effects as reflected by bone density show no significant differences between CD-1[ΔmGH.hGH] and CD-1[mGH] mice. However, CD-1[ΔmGH.hGH] mice show clear evidence of adipose tissue dysfunction, which can be exacerbated in mice fed a HFD. Finally, the potential for lactogenic activity to affect maintenance of glucose homeostasis in CD-1[ΔmGH.hGH] mice, directly or indirectly related to hGH, was observed. This potential to influence cells, tissues and/or processes or the response to stresses in addition to HFD cannot be ruled out. Thus, further assessment of other tissues and processes in males, as well as full assessments in female CD-1[ΔmGH.hGH] mice are warranted.

## Acknowledgements

This work was supported by a grant from the Canadian Institutes of Health Research (FRN-166215; PAC) and NSERC Discovery grant (RGPIN 05880-19; CGT). PAC holds the H.G. Friesen Chair in Endocrine and Metabolic Disorders.

## REFERENCES

[1] S.J. Frank, Growth hormone signalling and its regulation: preventing too much of a good thing, Growth Horm IGF Res 11(4) (2001) 201–12.

[2] J.M. Segrestaa, J. Gueris, M. Lamotte, [Current data on growth hormone], Pathol Biol (Paris) 23(5) (1975) 395–408.

[3] J.L. Kostyo, C.R. Reagan, The biology of growth hormone, Pharmacol Ther B 2(3) (1976) 591–604.

[4] R.E. Reagan, G.B. Pardue, E.J. Eisen, Predicting selection response for growth of channel catfish, J Hered 67(1) (1976) 49–53.

[5] A.J. Brooks, M.J. Waters, The growth hormone receptor: mechanism of activation and clinical implications, Nat Rev Endocrinol 6(9) (2010) 515–25.

[6] Y. Gan, A. Buckels, Y. Liu, Y. Zhang, A.J. Paterson, J. Jiang, K.R. Zinn, S.J. Frank, Human GH receptor-IGF-1 receptor interaction: implications for GH signaling, Mol Endocrinol 28(11) (2014) 1841–54.

[7] D.E. Berryman, E.O. List, Growth Hormone’s Effect on Adipose Tissue: Quality versus Quantity, Int J Mol Sci 18(8) (2017).

[8] N. Moller, J.O. Jorgensen, J. Moller, L. Orskov, P. Ovesen, O. Schmitz, J.S. Christiansen, H. Orskov, Metabolic effects of growth hormone in humans, Metabolism 44(10 Suppl 4) (1995) 33–6.

[9] J.J. Kopchick, D.E. Berryman, V. Puri, K.Y. Lee, J.O.L. Jorgensen, The effects of growth hormone on adipose tissue: old observations, new mechanisms, Nat Rev Endocrinol 16(3) (2020) 135–146.

[10] A. Bartke, J.J. Kopchick, The forgotten lactogenic activity of growth hormone: important implications for rodent studies, Endocrinology 156(5) (2015) 1620–2.

[11] P.A. Cattini, M.E. Bock, Y. Jin, J.A. Zanghi, H. Vakili, A useful model to compare human and mouse growth hormone gene chromosomal structure, expression and regulation, and immune tolerance of human growth hormone analogues, Growth Horm IGF Res 42-43 (2018) 58–65.

[12] T.C. Brelje, D.W. Scharp, P.E. Lacy, L. Ogren, F. Talamantes, M. Robertson, H.G. Friesen, R.L. Sorenson, Effect of homologous placental lactogens, prolactins, and growth hormones on islet B-cell division and insulin secretion in rat, mouse, and human islets: implication for placental lactogen regulation of islet function during pregnancy, Endocrinology 132(2) (1993) 879–87.

[13] J.L. Liu, K.T. Coschigano, K. Robertson, M. Lipsett, Y. Guo, J.J. Kopchick, U. Kumar, Y.L. Liu, Disruption of growth hormone receptor gene causes diminished pancreatic islet size and increased insulin sensitivity in mice, Am J Physiol Endocrinol Metab 287(3) (2004) E405–13.

[14] M. Freemark, I. Avril, D. Fleenor, P. Driscoll, A. Petro, E. Opara, W. Kendall, J. Oden, S. Bridges, N. Binart, B. Breant, P.A. Kelly, Targeted deletion of the PRL receptor: effects on islet development, insulin production, and glucose tolerance, Endocrinology 143(4) (2002) 1378–85.

[15] K. Lindberg, S.G. Ronn, D. Tornehave, H. Richter, J.A. Hansen, J. Romer, M. Jackerott, N. Billestrup, Regulation of pancreatic beta-cell mass and proliferation by SOCS-3, J Mol Endocrinol 35(2) (2005) 231–43.

[16] T.C. Brelje, L.E. Stout, N.V. Bhagroo, R.L. Sorenson, Distinctive roles for prolactin and growth hormone in the activation of signal transducer and activator of transcription 5 in pancreatic islets of langerhans, Endocrinology 145(9) (2004) 4162–75.

[17] D.W. Cooke, S.A. Sara, A. Divall, S. Radovick, Normal and Aberrant Growth in Children, in: S. Melmed, K.S. Polonsky, P.R. Larsen, H.M. Kronenberg (Eds.), Williams Textbook of Endocrinology, Elsevier 2016, pp. 964–1073.

[18] J. Auffret, M. Freemark, N. Carre, Y. Mathieu, C. Tourrel-Cuzin, M. Lombes, J. Movassat, N. Binart, Defective prolactin signaling impairs pancreatic beta-cell development during the perinatal period, Am J Physiol Endocrinol Metab 305(10) (2013) E1309–18.

[19] J.M.P. Holly, K. Biernacka, C.M. Perks, The Neglected Insulin: IGF-II, a Metabolic Regulator with Implications for Diabetes, Obesity, and Cancer, Cells 8(10) (2019).

[20] H. Modi, C. Jacovetti, D. Tarussio, S. Metref, O.D. Madsen, F.P. Zhang, P. Rantakari, M. Poutanen, S. Nef, T. Gorman, R. Regazzi, B. Thorens, Autocrine Action of IGF2 Regulates Adult beta-Cell Mass and Function, Diabetes 64(12) (2015) 4148–57.

[21] J.L. Kuk, P.T. Katzmarzyk, M.Z. Nichaman, T.S. Church, S.N. Blair, R. Ross, Visceral fat is an independent predictor of all-cause mortality in men, Obesity (Silver Spring) 14(2) (2006) 336–41.

[22] C. Pico, M. Palou, C.A. Pomar, A.M. Rodriguez, A. Palou, Leptin as a key regulator of the adipose organ, Rev Endocr Metab Disord 23(1) (2022) 13–30.

[23] Y. Zhang, K.Y. Guo, P.A. Diaz, M. Heo, R.L. Leibel, Determinants of leptin gene expression in fat depots of lean mice, Am J Physiol Regul Integr Comp Physiol 282(1) (2002) R226–34.

[24] M. Garaulet, N. Viguerie, S. Porubsky, E. Klimcakova, K. Clement, D. Langin, V. Stich, Adiponectin gene expression and plasma values in obese women during very-low-calorie diet. Relationship with cardiovascular risk factors and insulin resistance, J Clin Endocrinol Metab 89(2) (2004) 756–60.

[25] S. Parida, S. Siddharth, D. Sharma, Adiponectin, Obesity, and Cancer: Clash of the Bigwigs in Health and Disease, Int J Mol Sci 20(10) (2019).

[26] V.M. Sharma, E.T. Vestergaard, N. Jessen, P. Kolind-Thomsen, B. Nellemann, T.S. Nielsen, M.H. Vendelbo, N. Moller, R. Sharma, K.Y. Lee, J.J. Kopchick, J.O.L. Jorgensen, V. Puri, Growth hormone acts along the PPARgamma-FSP27 axis to stimulate lipolysis in human adipocytes, Am J Physiol Endocrinol Metab 316(1) (2019) E34–E42.

[27] Y. Jin, S.Y. Lu, A. Fresnoza, K.A. Detillieux, M.L. Duckworth, P.A. Cattini, Differential placental hormone gene expression during pregnancy in a transgenic mouse containing the human growth hormone/chorionic somatomammotropin locus, Placenta 30(3) (2009) 226–35.

[28] H. Vakili, Y. Jin, P.A. Cattini, Negative regulation of human growth hormone gene expression by insulin is dependent on hypoxia-inducible factor binding in primary non-tumor pituitary cells, J Biol Chem 287(40) (2012) 33282–92.

[29] H. Vakili, Y. Jin, J.I. Nagy, P.A. Cattini, Transgenic mice expressing the human growth hormone gene provide a model system to study human growth hormone synthesis and secretion in non-tumor-derived pituitary cells: differential effects of dexamethasone and thyroid hormone, Mol Cell Endocrinol 345(1-2) (2011) 48–57.

[30] H. Vakili, Y. Jin, P.A. Cattini, Energy homeostasis targets chromosomal reconfiguration of the human GH1 locus, J Clin Invest 124(11) (2014) 5002–12.

[31] H. Vakili, Y. Jin, P.A. Cattini, Evidence for a Circadian Effect on the Reduction of Human Growth Hormone Gene Expression in Response to Excess Caloric Intake, J Biol Chem 291(26) (2016) 13823–33.

[32] Y. Jin, J.S. Jarmasz, P.A. Cattini, Dexamethasone Rescues an Acute High-Fat Diet-Induced Decrease in Human Growth Hormone Gene Expression in Male Partially Humanized CD-1 Mice, DNA Cell Biol 40(3) (2021) 543–552.

[33] J.S. Jarmasz, Y. Jin, H. Vakili, P.A. Cattini, Sleep deprivation and diet affect human GH gene expression in transgenic mice in vivo, Endocr Connect 9(12) (2020) 1135–1147.

[34] E.O. List, D.E. Berryman, M. Buchman, E.A. Jensen, K. Funk, S. Duran-Ortiz, Y. Qian, J.A. Young, J. Slyby, S. McKenna, J.J. Kopchick, GH Knockout Mice Have Increased Subcutaneous Adipose Tissue With Decreased Fibrosis and Enhanced Insulin Sensitivity, Endocrinology 160(7) (2019) 1743–1756.

[35] J.R. Eveleigh, H.H. Jasdrow, M. Gunner, The breeding performance of CD1 stud male mice with some comparative data from BALB/c, PSD and PSDI strains, Lab Anim 17(3) (1983) 230–4.

[36] C.N. Peroni, C.Y. Hayashida, N. Nascimento, V.C. Longuini, R.A. Toledo, P. Bartolini, C.Y. Bowers, S.P. Toledo, Growth hormone response to growth hormone-releasing peptide-2 in growth hormone-deficient little mice, Clinics (Sao Paulo) 67(3) (2012) 265–72.

[37] H. Vakili, Jin, Y., Menticoglou, S., Cattini, P. A., CCAAT-enhancer-binding protein beta (C/EBPbeta) and downstream human placental growth hormone genes are targets for dysregulation in pregnancies complicated by maternal obesity, Journal of Biological Chemistry 288(31) (2013) 22849–61.

[38] S. Moazzam, N. Noorjahan, Y. Jin, J.I. Nagy, E. Kardami, P.A. Cattini, Effect of high fat diet on maternal behavior, brain-derived neurotrophic factor and neural stem cell proliferation in mice expressing human placental lactogen during pregnancy, J Neuroendocrinol (2023) e13258.

[39] C.A. Schneider, W.S. Rasband, K.W. Eliceiri, NIH Image to ImageJ: 25 years of image analysis, Nat Methods 9(7) (2012) 671–5.

[40] V. DeClercq, P. Zahradka, C.G. Taylor, Dietary t10,c12-CLA but not c9,t11 CLA reduces adipocyte size in the absence of changes in the adipose renin-angiotensin system in fa/fa Zucker rats, Lipids 45(11) (2010) 1025–33.

[41] F. Nassir, R.S. Rector, G.M. Hammoud, J.A. Ibdah, Pathogenesis and Prevention of Hepatic Steatosis, Gastroenterol Hepatol (N Y) 11(3) (2015) 167–75.

[42] T.L. Stanley, S.K. Grinspoon, Effects of growth hormone-releasing hormone on visceral fat, metabolic, and cardiovascular indices in human studies, Growth Horm IGF Res 25(2) (2015) 59–65.

[43] W.Y. Chen, D.C. Wight, T.E. Wagner, J.J. Kopchick, Expression of a mutated bovine growth hormone gene suppresses growth of transgenic mice, Proc Natl Acad Sci U S A 87(13) (1990) 5061–5.

[44] Y. Qian, D.E. Berryman, R. Basu, E.O. List, S. Okada, J.A. Young, E.A. Jensen, S.R.C. Bell, P. Kulkarni, S. Duran-Ortiz, P. Mora-Criollo, S.C. Mathes, A.L. Brittain, M. Buchman, E. Davis, K.R. Funk, J. Bogart, D. Ibarra, I. Mendez-Gibson, J. Slyby, J. Terry, J.J. Kopchick, Mice with gene alterations in the GH and IGF family, Pituitary 25(1) (2022) 1–51.

[45] S. Lind, M. Rudling, S. Ericsson, H. Olivecrona, M. Eriksson, B. Borgstrom, G. Eggertsen, L. Berglund, B. Angelin, Growth hormone induces low-density lipoprotein clearance but not bile acid synthesis in humans, Arterioscler Thromb Vasc Biol 24(2) (2004) 349–56.

[46] Y. Fan, R.K. Menon, P. Cohen, D. Hwang, T. Clemens, D.J. DiGirolamo, J.J. Kopchick, D. Le Roith, M. Trucco, M.A. Sperling, Liver-specific deletion of the growth hormone receptor reveals essential role of growth hormone signaling in hepatic lipid metabolism, J Biol Chem 284(30) (2009) 19937–44.

[47] B. Moreno-Carranza, M. Bravo-Manriquez, A. Baez, M.G. Ledesma-Colunga, X. Ruiz-Herrera, P. Reyes-Ortega, E.A. de Los Rios, Y. Macotela, G. Martinez de la Escalera, C. Clapp, Prolactin regulates liver growth during postnatal development in mice, Am J Physiol Regul Integr Comp Physiol 314(6) (2018) R902–R908.

[48] N.C. Olarescu, K. Gunawardane, T.K. Hansen, N. Moller, J.O.L. Jorgensen, Normal Physiology of Growth Hormone in Adults, in: K.R. Feingold, B. Anawalt, M.R. Blackman, A. Boyce, G. Chrousos, E. Corpas, W.W. de Herder, K. Dhatariya, K. Dungan, J. Hofland, S. Kalra, G. Kaltsas, N. Kapoor, C. Koch, P. Kopp, M. Korbonits, C.S. Kovacs, W. Kuohung, B. Laferrere, M. Levy, E.A. McGee, R. McLachlan, M. New, J. Purnell, R. Sahay, A.S. Shah, F. Singer, M.A. Sperling, C.A. Stratakis, D.L. Trence, D.P. Wilson (Eds.), Endotext, South Dartmouth (MA), 2000.

[49] A.L. Castell, C. Goubault, M. Ethier, G. Fergusson, C. Tremblay, M. Baltz, D. Dal Soglio, J. Ghislain, V. Poitout, beta Cell mass expansion during puberty involves serotonin signaling and determines glucose homeostasis in adulthood, JCI Insight 7(21) (2022).

[50] A. Villaret, J. Galitzky, P. Decaunes, D. Esteve, M.A. Marques, C. Sengenes, P. Chiotasso, T. Tchkonia, M. Lafontan, J.L. Kirkland, A. Bouloumie, Adipose tissue endothelial cells from obese human subjects: differences among depots in angiogenic, metabolic, and inflammatory gene expression and cellular senescence, Diabetes 59(11) (2010) 2755–63.

[51] C.M. Gorvin, The prolactin receptor: Diverse and emerging roles in pathophysiology, J Clin Transl Endocrinol 2(3) (2015) 85–91.

[52] C. Ling, G. Hellgren, M. Gebre-Medhin, K. Dillner, H. Wennbo, B. Carlsson, H. Billig, Prolactin (PRL) receptor gene expression in mouse adipose tissue: increases during lactation and in PRL-transgenic mice, Endocrinology 141(10) (2000) 3564–72.

